# Label Noise Limits TCR-pMHC Specificity Prediction: Improved Performance Through AlphaFold3-Based Structural Modeling and Data Denoising

**DOI:** 10.64898/2026.08.25.746957

**Authors:** Pilar Ballesteros-Cuartero, Johanne Lund, Morten Nielsen

## Abstract

T cell receptor (TCR) binding to peptides presented by major histocompatibility complex (MHC) molecules is a key step in T cell activation, and forms the basis of adaptive immunity. Predicting this specificity is therefore essential to developing effective TCR-based immunotherapies and vaccines. Despite its clinical relevance, predicting TCR-pMHC specificity for previously unseen peptides remains an open problem, with structural modeling so far the only strategy showing any predictive power in this setting. In this study, we find that this limited performance is substantially driven by label noise in the data used to train and evaluate these methods, an effect that has so far been largely underexplored. Using an AlphaFold3-based pipeline adapted for TCR-pMHC structural modeling, we achieve state-of-the-art specificity prediction, outperforming AlphaFold2.3-based and sequence based methods, and performing at par with the leading Immrep2025 competition submission. Combining this pipeline with a cluster-based denoising algorithm, we show that removing mislabeled points from a large specificity dataset increased binder ranking accuracy by more than 70% relative to the full dataset. Together, these results highlight label noise as a major factor limiting the performance that any method in this field can achieve, and show that combining structural modeling with label denoising substantially improves TCR-pMHC specificity prediction, making such approaches an attractive complement to current sequence-based approaches for refining TCR target selection.

## Introduction

T cells are essential for identifying and eliminating infected or malignant cells ^1^. This recognition is mediated by the T cell receptor (TCR), which binds to peptides presented on the cell surface by major histocompatibility complex (MHC) molecules. Accurately characterizing the specificity and cognate peptide-MHC targets of a given TCR would hence directly accelerate the development of TCR-based immunotherapies and vaccines ^2–5^.

Given this, several methods have been developed to predict TCR-pMHC specificity. Current approaches can predict binding to peptides seen during training ^6,7^, but generalizing to unseen epitopes remains a challenge ^8^, despite this setting being crucial to TCR-based therapy development. In previously reported benchmarks, sequence based prediction methods, using either machine learning or distance based approaches, have shown to perform near random discrimination in this unseen epitope setting ^8^. In contrast, structure based methods have shown some predictive power here, since they can capture general binding patterns shared across TCRs at the structural level, without relying directly on sequence similarity to peptides seen during training ^8,9^.

Given the specific traits of the TCR-pMHC system, such as the lack of co-evolution across chains and the high flexibility of the TCR complementarity determining regions (CDRs), methods tailored specifically to TCR-pMHC structure modeling have been demonstrated to outperform general-purpose structural tools ^10^. Some of these methods focus on ensuring the correct docking of TCR-pMHCs ^9^, others on the creation of curated databases and joint pMHC modeling ^11^, while others combine structure and sequence data in multimodal deep learning frameworks ^12–18^. However, even with these tailored methods, specificity prediction for unseen epitopes remains a challenge ^8^.

Beyond these modeling challenges specific to TCR-pMHC, another critical yet less studied element could also be limiting predictive accuracy. Public TCR-pMHC specificity datasets are known to contain a significant amount of mislabeled entries, with some reported points being falsely labeled as positives ^19^. This suggests that the predictive performance of current models may be underestimated, since part of this modest predictive power could reflect data noise rather than modeling limitations. Further, this aspect places a cap on developments within the field, since the substantial noise level introduces artifacts that may interfere with learnability in the data. Despite this, the effect of label noise on model training and evaluation in TCR-pMHC specificity prediction remains largely unexplored.

Here, we present NetTCRfold, an AlphaFold3-based ^20^ pipeline adapted for TCR-pMHC structural modeling, to address these questions and challenges. We first investigate how the essential building blocks of AlphaFold3, such as MSA construction and template selection, can be refined in the context of TCR-pMHC complexes, and next benchmark the resulting pipeline to validate modeling and TCR-pMHC docking performance. We also investigate which AF3-derived metrics perform best for model selection and specificity prediction. We then evaluate the pipeline’s specificity prediction performance against state-of-the-art methods on the Immrep2025 competition dataset, and further assess its performance on a larger, sequence-based dataset derived from Deleuran et al. ^21^. Finally, we apply a clustering-based denoising algorithm, and compare predictive performance of our pipeline before and after denoising to quantify the effect of label noise.

## Materials and Methods

### NetTCRfold pipeline

The structural modeling pipeline was developed by introducing a set of modifications to the AlphaFold3 (AF3) ^20^ code and its associated databases, as described in detail below.

#### Reduction of the Multiple Sequence Alignment Database

We used the TCRModel2^11^ sequence databases, which consist of reduced versions of UniRef90^22^, UniProt^23^ and BFD^24^ containing only TCR and MHC related sequences. The authors excluded the metagenomic database^25^, as no hits were found for TCR-pMHC complexes. The resulting MSA databases contain 52,096 TCR-related sequences and 137,991 MHC-related sequences, and each database contributes as follows: 450 sequences for Small BFD, 43,638 for UniRef90 and 145,999 Uniprot sequences.

#### Multiple Sequence Alignment Modification

The MSA generation step in AlphaFold3 has two different components: an unpaired alignment and a paired alignment. MSA is first performed independently for each chain against each database. In the case of paired alignments, sequences from the same organism, as annotated by UniProt, are then paired row-wise across chains to capture cross-chain coevolution, with the pairing order determined by query similarity. UniProt derived sequences that could not be matched across chains, as well as those derived from BFD or UniRef90, are retained in a diagonal sparse matrix, which is what we call unpaired MSA. In the case of only using unpaired alignments, only the hits from the BFD or UniRef90 databases are included in the diagonal sparse matrix.

By default, AF3 accepts only two MSA configurations, either no-MSA or full-MSA, and using only paired or unpaired MSA is not supported. Therefore, in this work, we modified the AF3 pipeline to allow any of the four different configurations: no MSA, unpaired-only, paired-only or full MSA. This was done by modifying the outputs of the data generation pipeline prior to inference, by keeping only the desired MSA.

#### TCR template search

AlphaFold3 constructs a hidden Markov model (HMM) from the unpaired MSA and performs a search against entries with known structure using this HMM profile. The four most similar matches to the HMM are then retrieved, with similarity determined by HMMER’s ^26^ score and E-value (higher score and lower E-value indicating higher similarity), and their structures are used as templates.

Prior work has shown that when the whole MSA is used for template search, immunoglobulins are frequently misidentified as TCR templates ^11^. To address this, the authors of TCRModel2 proposed restricting template search to the query sequence alone. We adopted this modification for both TCR and MHC chains.

#### Template database reduction

By default, AF3 searches the entire RCSB Protein Data Bank for templates (https://www.rcsb.org/)^27^. This database contains a large proportion of structures unrelated to TCR-pMHC biology, which both dilutes template relevance and increases search time. We therefore curated a reduced, TCR and MHC specific template database. We first compiled a set of relevant TCR and MHC entries from STCRDab^28^ and IEDB^29^, and then identified which of these entries were present in the RCSB sequence records database, the database AF3 uses by default to perform template search.

In the first step, we downloaded the full STCRDab database, which contains only TCR-pMHC complexes, together with the IEDB database for pMHC complexes (accessed on 17 February 2026). STCRDab entries were filtered according to the following criteria: TCR type = αβTCR, antigen type = peptide, MHC type = MHC class I or II. The pMHC IEDB database was filtered to MHC Class I or II, or non-classical MHC restricted to the alleles HLA-E*01:03, HLA-G*01:01, HLA-E*01:01, HLA-G, H2-Q9, H2-M3, H2-Qa-1a, and H2-Qa-1b. Entries lacking a chain assignment for any component were excluded, since AF3 requires a specific PDB ID and chain identifier to retrieve the templates, as were entries in which the antigen and the MHC molecule were assigned to the same chain. This curation resulted in 2,939 unique MHC chains, 443 to TCRα chains and 443 to TCRβ chains, a total of 3,825 unique sequences.

In the second step, we cross-referenced these 3,825 sequences against the RCSB PDB sequence database provided by the TCRmodel2 authors, and found that 3,051 were present in this database (see Supplementary data S1 for the excluded entries). This two-step process reduced AF3’s template search space from 667,140 sequences in the full database to the 3,051 curated, TCR and MHC relevant sequences introduced here (see Supplementary Data S2 for entry-level information on the final curated database).

#### Modeling pipeline

In the final modeling pipeline, AF3 models were generated by using unpaired MSAs computed for both TCR chains and the MHC chain. Template search was restricted to the query sequence for the MHC and both TCR chains. The peptide was excluded from both the MSA and template search steps due to its short length. For each datapoint, 50 models were generated using 10 seeds and 5 diffusion samples per seed. The top-ranked model for each datapoint was subsequently selected by max-pooling a chosen confidence (see below) metric across all 50 models, a process we refer to as model selection throughout this work.

#### Model scoring

For each model, we extract a large series of AF3-derived-confidence metrics, including predicted global Template Modelling (TM) score (pTM, an estimate of the complex overall structural accuracy), predicted interface TM-score (ipTM, a variant of pTM restricted to chain interfaces and combined across said interfaces), and the composite AF3 confidence score (0.2 x pTM + 0.8 x ipTM). We additionally extracted chain-specific and intra-chain pairwise metrics, including per-chain pTMs, per-chain ipTMs, pairwise chain-chain ipTM, and pairwise chain-chain predicted aligned error (PAE). Because PAE values are inversely related to model confidence (lower values indicate higher confidence), we transformed PAE into a bounded confidence on a [0,1] scale using the formula 1. The score was calibrated such that a PAE of 10 Å maps to a confidence score of 0.5, a PAE of 0 Å maps to a score of 1, and scores below 0.5 correspond to progressively higher PAE values.

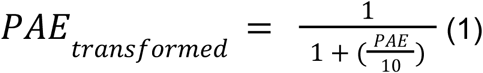

We further extracted the mean predicted local difference distance test (pLDDT) scores across the whole complex normalized to a score of [0,1], as well as a pLDDT score restricted to the peptide and to the complementarity-determining regions (CDRs). Mean PAE was also computed for specific CDR-chain pairings, namely CDR1/CDR2 with the MHC chain and CDR3 with the peptide. The presence of an interface between any two chains was evaluated, with an interface defined as the presence of at least one pair of heavy atoms within 5 Å of one another. Finally, we derived the interaction prediction Score from Aligned Errors (ipSAE)^30^, the ipSAE_d0chn, which is a variant of the ipSAE where the scaling factor (d0) is set to the sum of the length of two chains under study, and the ipSAE_d0dom, where the d0 factor is the sum of residues with a PAE lower than 10 Å.

PAE and iPSAE are asymmetric metrics, meaning that their value depends on the order in which the two chains are considered, for example, TCRα-peptide PAE differs from peptide-TCRα PAE. To obtain a single, symmetric value for each chain pair, we took the maximum of these two scores (Formula 2). Independently, for metrics computed separately for TCRα and TCRβ, we derived a single composite TCR-level metric by taking the maximum value across the two chains (Formula 3). For datapoints with a corresponding solved structure, DockQ version 1.0 ^31^ was additionally computed for the TCR-pMHC interface.

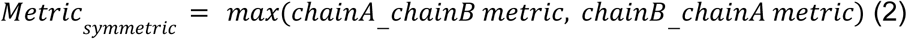

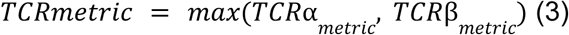

### Benchmarking dataset of solved structures

The benchmarking dataset for pipeline and metric selection was assembled from 38 solved αβTCR-pMHC class I structures, combining 25 structures from the benchmark dataset introduced by Deleuran et al. ^21^ with 13 additional structures released after NetTCR-struc’s benchmark curation (Supplementary Data S3). Briefly, all the structures were retrieved from the RCSB PDB and restricted to those released after the AlphaFold3 training data cutoff (30 September 2021), with a resolution cutoff of 3.5 Å. Complexes containing non-standard amino acids in the peptide were excluded. For all retained complexes, only the α chain of the MHC was used, and the TCRs were trimmed to their variable domains using the ANARCI^32^ *run_anarci* function (version 2024.05.21). The complexes were then redundancy reduced using the Hobohm 1 algorithm with a 95% sequence similarity threshold, meaning that complexes with either a TCRa or TCRb similarity higher than this threshold to a datapoint previously encountered were removed from the benchmark dataset. An additional filtering step was included to remove any benchmark structure whose TCRα or TCRβ sequence was more than 95% similar to any structure released before the AF3 training data cutoff.

### Template diversity quantification

Template diversity was quantified per chain and per datapoint as the number of unique templates selected for the given query. Here, a template was identified by its PDB identifier and chain ID.

### Structural metrics

Structural metrics were computed by comparing the AF3-generated models with their corresponding structures, using Cα atoms only to assess the overall backbone accuracy. All metrics were computed using the Python package biotite ^33,34^ (version 1.5.0). The global TM-score was obtained by superimposing the model and the template structures with superimpose(), followed by tm_score() ^35^. The TCR-peptide interface TM-score (iTM_tcr_pep) was obtained by first defining the TCR-peptide interface as any pair of Cα atoms within 14 Å of one another and then restricting the TM-score calculation to these interface atoms. Per-chain lDDT scores were computed by passing the model and reference chains to lddt(). Per-chain Root Mean Square Deviation (RMSD) was computed after aligning the model and the reference structures on the MHC chain, which served as a structural anchor given its high conservation and generally accurate modelling. The alignment was performed using superimpose() with an atom mask restricted to MHC atoms in the reference structure. Following MHC-based alignment, RMSD was computed per chain using rmsd(). This metric is referred to as alignedChainRMSD, where “Chain” denotes the specific chain under evaluation.

### Structure alignment visualization and model displacement quantification

Structures were visualized in PyMOL (version PyMOL 3.0.0 Open-Source, 2024-05-27). The model and the reference structures were aligned on the MHC chain, consistent with the RMSD calculation described above. The reference MHC chain was selected using cmd.select(“ref_chain”, “reference_structure and chain A”), and each model was aligned to this reference using [cmd.align(f”{obj} and chain A”, “ref_chain”) for obj in cmd.get_object_list() if obj != “reference_structure”].

Per-residue displacement between model and reference CDRs, following MHC-based superimposition, was defined as the Euclidean distance between the corresponding Cα atoms. CDR indices for TCRa and TCRb were obtained using IMGT numbering scheme with ANARCI *number* function (version 2024.05.21), and displacement was obtained for the CDR residues only.

### Selection of a metric for model retrieval from model pool

For each datapoint, the modeling pipeline generates 50 candidate models, from which the model best representing the underlying structure needs to be identified (see Modeling pipeline and Model scoring for more details). For the benchmark dataset, where solved structures are available, DockQ was used as the reference metric for model selection. For each datapoint, the model with the maximum DockQ score was retrieved, a selection strategy we refer to as the Oracle throughout this work.

In the absence of a solved structure, model selection must rely on an AF3-derived confidence metric. To identify which metric best approximates Oracle-based selection, we first removed from the candidate pool any model lacking a predicted TCRα-peptide or TCRβ-peptide interface. The top-ranked model for each datapoint was then identified by max-pooling a given confidence metric, and the corresponding DockQ score was retrieved. For data points where all 50 models were excluded during the filtering step, a random model was selected.

### Batch construction and metrics for target selection

As stated above, in this work, we use the term target selection to refer to the process of identifying the true binder within a pool of candidate TCR-pMHC complexes that also includes non-binders.

For all the cases where target-selection performance is assessed, we adopted the batch setup introduced by Deleuran et al. [21]. Here, for each pMHC in a positive TCR-pMHC complex, negatives were generated by swapping the TCR with 5 TCRs positive to other pMHCs. For a TCR to be eligible for the swap, the peptide pairs had to have a Levenshtein distance bigger than 3. Each positive complex, together with its five swapped negatives, constitutes one batch.

Next, for each batch, we evaluated the rank of the true positive relative to the five negatives, using a metric introduced by Delauran et al. ^21^, called the true-positive rank metric (TPR). This metric captures how many non-binding complexes are scored higher than the true positive and it is obtained as follows (Formula 4):

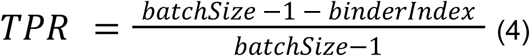

Here, binderIndex is the rank of the true positive within the batch, with all complexes ranked according to a given confidence metric and rank 0 assigned to the highest-scoring complex. In cases of ties the binderIndex is defined as the mean rank of all tied complexes.

To track changes in TPR across batches, we defined a cumulative TPR, computed by first ranking batches from highest to lowest binding-metric score, and then summing TPF cumulatively across the ranked batches. Cumulative TPR was computed starting from the sixth batch onward, to avoid instability arising from small batch numbers early in the ranking. Performance was quantified as the area under the cumulative TPR curve (AUC).

### Target selection benchmark using the solved-structure dataset

A target-selection benchmark was then created from the solved-structure benchmark introduced above, in order to identify the best metrics for target-selection. Using the batch construction procedure described above, five negatives were generated for each positive TCR-pMHC by swapping its TCR with five TCRs binding to different pMHC. This resulted in 38 batches, each with a 1:5 positive-to-negative ratio, comprising 38 positive and 190 negative examples in total. Each complex was modeled with 10 seeds and 5 diffusion samples per seed, giving 50 models per TCR-pMHC complex. The best model per complex was selected using one four candidate metrics: MHC-CDR1/2 PAE mean metric, TCR-pep ipSAE d0dom metric, AF3 confidence metric and a NetTCRstruc derived metric (NetTCRstruc-IF1-AF3confidence).

This batch setup was used to identify the metric best suited for discriminating binders from non-binders (metric for target-selection). We evaluated each metric’s discrimination performance by computing the True-Positive Rank metric for each batch, and reported the mean across all batches.

### Immrep2025 data modeling and evaluation

The Immrep2025^8^ dataset consists of 10,000 TCR-pMHC complexes spanning two HLA alleles (HLA-A02:01 and HLA-B40:01) and 20 peptides. All complexes were modeled using our NetTCRfold pipeline, generating 50 models per complex. Model selection per datapoint was done using either the mean PAE between MHC and CDR1/2 residues (MHC-CDR1/2 PAE mean) or the TCR-peptide ipSAE d0dom metric (for details on these metrics see model selection), and target selection was performed using TCR-peptide ipSAE d0dom metric in both cases.

For method evaluation, we assessed the performance of our method on the private IMMREP25 dataset, as defined by the Immrep2025 organizers, to ensure comparability with the competition’s published results. The private dataset was derived as follows. We first subsetted the full dataset to the 8,938 records randomly selected by the organizers. Each record was then classified into a public or private subset according to the criteria described by the authors (2.2 Evaluation dataset processing in Immrep2025 manuscript) ^8^. This resulted in a private dataset of 7,344 records, spanning the 18 private peptides and both HLA-alleles, matching the dataset size reported in the original publication. Per-peptide AUC0.1 was then computed for each of the 18 private peptides using the TCR-peptide ipSAE d0dom metric as target selection metric. Per-peptide AUC0.1 for the top-performing competition submissions were obtained from Supplementary Figure 3 from the Immrep25 competition report.

### TCR specificity prediction

The sequence-based TCR specificity prediction dataset was obtained from Deleuran et al. ^21,36^. Shortly, the authors obtained positive examples from IEDB^29^, VDJdb^37^ and a 10X Genomics dataset denoised using the ITRAP algorithm^38^. The dataset was downsampled to a maximum of 200 positive examples per peptide, and negative examples were generated using the swapping approach described above, with each positive TCR from a specific TCR-pMHC complex swapped by 5 TCRs positive to other pMHCs. The resulting dataset comprised 2,945 positive examples and 14,725 swapped negative examples, giving a total of 17,670 datapoints.

The full dataset was modeled using our NetTCRfold pipeline, generating 50 models per complex. The best model for each datapoint was selected using the MHC-CDR1/2 PAE mean metric and the TCR-pep ipSAE d0dom metric was subsequently used for specificity prediction. Corresponding AF2 derived models and metrics were obtained directly from the authors of NetTCR-struc ^21^.

### TCRvdb

The TCR specificity prediction dataset defined above was subset to the TCR-pMHC complexes evaluated in Messemaker et al. denoising study ^19^ and available in TCRvdb. This resulted in a dataset containing 96 TCR-pMHC complexes. Of the 54 entries from YLQPRTFLL, 35 were confirmed as true binders and 19 as noise (false positives). Of the 42 GLCTLVAML related complexes, 28 were confirmed as true binders and 14 as false positives. The final denoised dataset therefore comprised 63 experimentally confirmed true binders and 33 experimentally confirmed non-binders (false positives).

We then retrieved, from the batch-sampling experiment, the batches corresponding to these 96 data points together with their associated binding metrics. Cumulative TPR was computed for the full 96-datapoint subset, and for the two subsets containing the true-binder and noise batches, respectively.

### Clustering-based denoising algorithm

The TCRdenoise denoising tool is a sequence-based method for separating binding and non-binding TCRs in a peptide-specific repertoire. It uses TCRbase ^39,40^ and TCRdist3 ^41,42^ to compute pairwise TCR distance matrices, from which clustering solutions are generated using agglomerative clustering across a range of distance thresholds. The clustering solutions are then evaluated using a refined silhouette score modified to handle singletons to identify the optimal distance threshold for each distance matrix. Next, TCRs located in clusters are classified as binders, and non-clustered TCRs as noise (non-binders). Finally, the selected clustering solutions obtained from the two similarity metrics are combined into a consensus clustering. Further details of the “denoising algorithm” are described in Supplementary Figure 7.

### Denoising the TCR specificity dataset

Positive complexes in the TCR specificity dataset were denoised using the sequence based clustering algorithm described above. Batches from the batch-sampling experiment were partitioned according to whether their associated positive complex was classified as noise or as a true binder. This resulted in 1,650 datapoints identified as binders and 1,295 as noise. Cumulative TPR and the corresponding area under the cumulative TPR curve were then computed separately for each of the two groups.

### Training of NetTCR

NetTCR^36^ is an ensemble of convolutional neural networks used for predicting TCR-pMHC binding. In short, the networks take as input the peptide and 6 CDR loop sequences. Each input sequence is passed through a CNN layer, and the max-pooled output fed into a dense layer and onward to the output neuron (for details refer to ^36^).

In this study, we retrain NetTCR on the full and the denoised TCR specificity data described above, respectively. The models were trained in a 5 fold nested cross-validation setup. As the original data already had redundancy reduced, each dataset was randomly split into five partitions, with the requirement that each partition had a minimum of 5 TCRs per peptide. This resulted in removing 8 peptides (FEDLRLLSF, FEDLRVLSF, GPRLGVRAT, RLPGVLPRA, RLRAEAQVK, RPPIFIRRL, SLFNTVATLY, and VLFGLGFAI) from the the denoised data reducing the total number of positive from 1,650 to 1,496 TCRs. Negative data were generated by swapping the TCRs for a given peptide with TCRs binding to other peptides. For each model, the performance evaluation was done in a per peptide manner in terms of the AUC0.1 value calculated from the concatenated test set predictions. For details on the data partitioning procedure, model training, model architecture, and performance evaluation refer to the original NetTCR-2.2 publication ^36^.

## Results

In this work, we investigate how a tailored version of AlphaFold3 can be used for accurate structural modeling of TCR-pMHC complexes and assessment of TCR specificity. The tailoring is made through modifications to the data pipeline and exploration of different metrics for model selection. Next, we evaluate the resulting pipeline’s ability to predict TCR-pMHC specificity across independent datasets. Finally, we examine the effect of noise in TCR specificity prediction, introduce a clustering-based algorithm for unsupervised denoising, and assess the impact of noise on specificity prediction performance.

### Adapting Alphafold3 for TCR-pMHC structure prediction

Alphafold3’s structure prediction pipeline relies on two data-generation steps, multiple sequence alignment (MSA) construction and per-chain structural template search. In the default pipeline, both are tailored to general purpose predictions rather than TCR-pMHC specific predictions. Therefore, in this work both steps were modified to better reflect the specific scenario of modelling TCR-pMHC complexes. We assess the impact of the changes on a benchmark dataset consisting of 38 TCR-pMHC class I solved structures, spanning 31 distinct peptides (for more details see methods). Each datapoint is modeled with 10 seeds and 5 diffusion samples, giving 50 models per datapoint (see methods for more details). The top model per datapoint is selected using DockQ as the reference metric, representing the ceiling performance of the pipeline, which we refer to as Oracle throughout this work. Next, we use different confidence metrics derived from the generated structure, including the AF confidence metric, to select the max-pooled top1 model from the 50 model pool to estimate the real performance of the model, in the absence of ground truth.

#### 1.1 Adapting the MSA search step for TCR-pMHC

##### A curated TCR-MHC specific database reduced MSA runtime by up to 51-fold with minimal accuracy cost

AF3’s default MSA step searches each query against four large sequence databases. Since the majority of these sequences are unrelated to TCRs and MHCs, this search step constitutes a major speed bottleneck without providing an increase in relevant information retrieval. To solve this speed bottleneck, we replaced the default databases with a curated, TCR-MHC restricted database as originally introduced in TCRmodel2 ^11^ (see methods).

This substitution reduced MSA construction time by approximately 17-fold per chain, corresponding to a 53-fold reduction in total MSA runtime per complex, going from 54 minutes per complex (i.e 18 minutes per chain x 3 chains) to 1 minute (Figure 1A). We tested the change in model accuracy when changing the MSA database while keeping the rest of the pipeline to the AF3 default, and saw that this speedup came at a small cost to model quality on our benchmark dataset (Figure 1B). However, fast modeling is required to make large-scale TCR-pMHC predictions more accessible, so we believe the change represents a favourable trade-off between runtime and accuracy.

**Figure 1:**
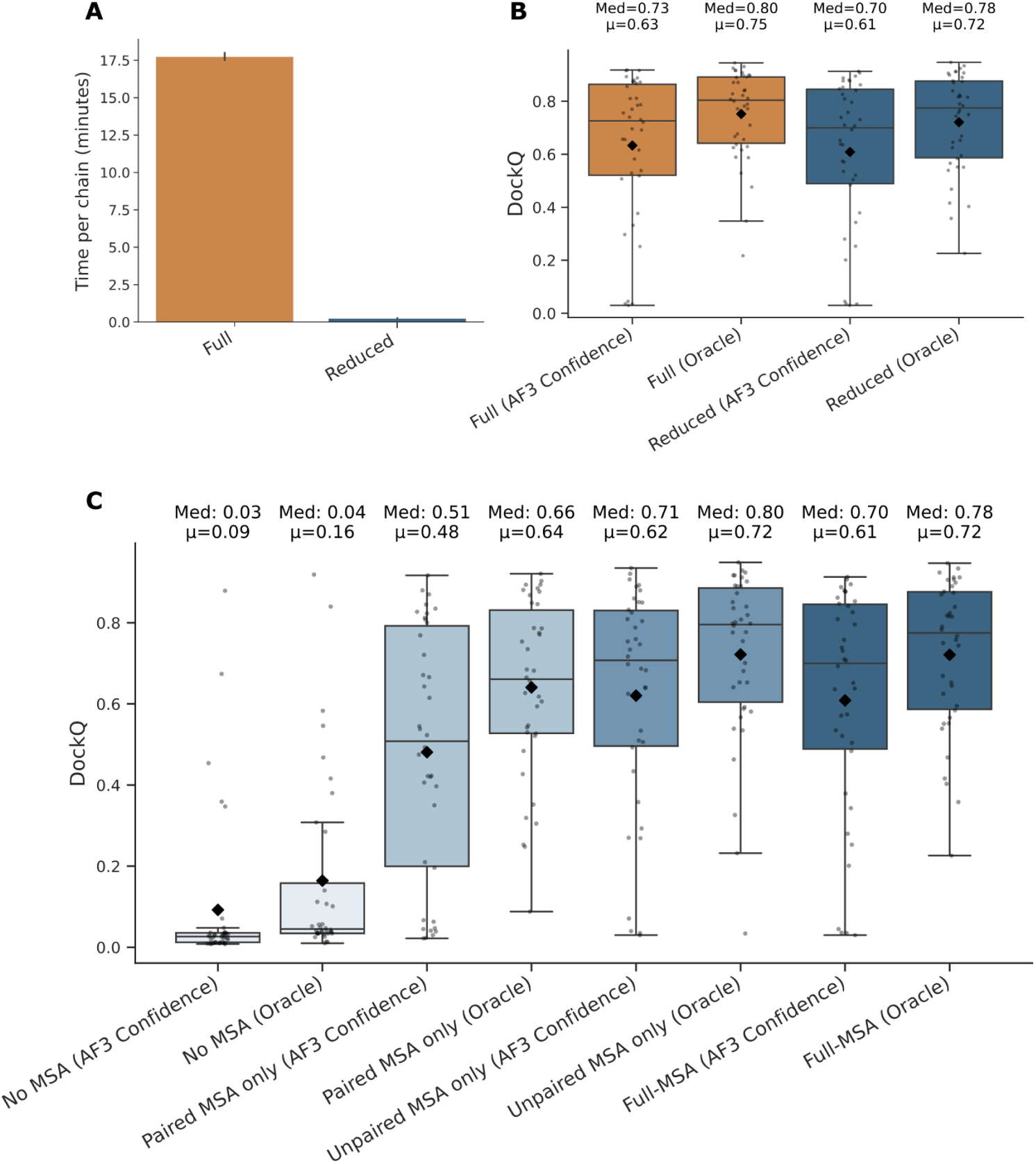
Impact of MSA database size and MSA pairing strategy on TCR-pMHC structure prediction. A) MSA construction time (minutes per chain) using the full sequence database versus a TCR/MHC restricted database. B) Effect of MSA database size on model quality across the benchmark dataset (n=38), using standard template selection. C) Effect of MSA pairing strategy on model quality, evaluated on the benchmark dataset (n=38) using the reduced MSA database and standard template selection. For B) and C), each datapoint was modeled with 10 seeds and 5 diffusion samples per seed (50 models per datapoint). The top model was selected by max-pooling across the 50 models, using either DockQ (Oracle) or AF3 confidence. Model quality is reported as the DockQ of the selected model according to either metric. Each point represents a TCR-pMHC complex, and the mean is shown by the diamond point.

##### Unpaired MSAs are sufficient for a system without cross-chain co-evolution

AF3 performs a cross-chain MSA pairing by species, a strategy designed to capture co-evolution between chains in a complex that have evolved together. However, this assumption does not hold for TCR-pMHC complexes, since in this case pairing TCRa and TCRb chains by species would fail to reflect the underlying biology behind TCR chain pairing. We therefore hypothesized that restricting the MSA to an unpaired configuration, would maintain or improve modelling accuracy be better reflecting this lack of co-evolution.

We tested several combinations of paired and unpaired MSA and found that the inclusion of MSAs is required for AF3 to model TCR-pMHC complexes correctly. Among the tested combinations, using unpaired MSA only improved modeling accuracy relative to the default AF3 scheme (Unpaired + Paired MSA), consistent with the biology of our system (Figure 1C). Based on this result, we adopted unpaired-only MSA throughout the remainder of the pipeline, and all subsequent results use this configuration.

#### 1.2 Adapting template search step for TCR-pMHC

##### Query-based template search combined with a curated template database increases template diversity and improves model quality

Previous work has suggested that restricting the template search to the chain sequence alone, rather than its full MSA profile, can improve TCR-pMHC modelling accuracy, since MSA-based search can lead to the identification of immunoglobulins as TCR templates ^11^. We evaluated this modification in our pipeline and examined its effect on template selection.

Using the full MSA profile for template search resulted in an extremely low template diversity, with the same single template (6ZQK_A) being selected for TCRβ across every datapoint, and only two unique templates selected for TCRα (6ZQK_A, 4HJJ_L). In these cases, different regions of the same structure were repeatedly selected as templates, rather than the search selecting a more diverse pool to fully exploit the four templates available per chain. Moreover, consistent with previous reports, both of these templates were immunoglobulins rather than TCR structures. Restricting the template search to the chain sequence alone substantially increased template diversity, giving an average of four unique templates per chain across the benchmark dataset. This increase was also reflected in the number of unique templates identified across the full dataset, with template identity varying between datapoints. The gain in diversity was most pronounced for the TCRα and TCRβ chains, where 73 and 85 unique templates were selected, respectively, out of a possible 152 (i.e four unique templates per datapoint across all 38 datapoints) (Figure 2A). Taken together, these results indicate that the template search modification shifts this step from being driven by a generic TCR MSA profile with datapoint-agnostic template selection to one driven by each datapoint’s unique sequence, resulting in datapoint-specific template selection.

**Figure 2:**
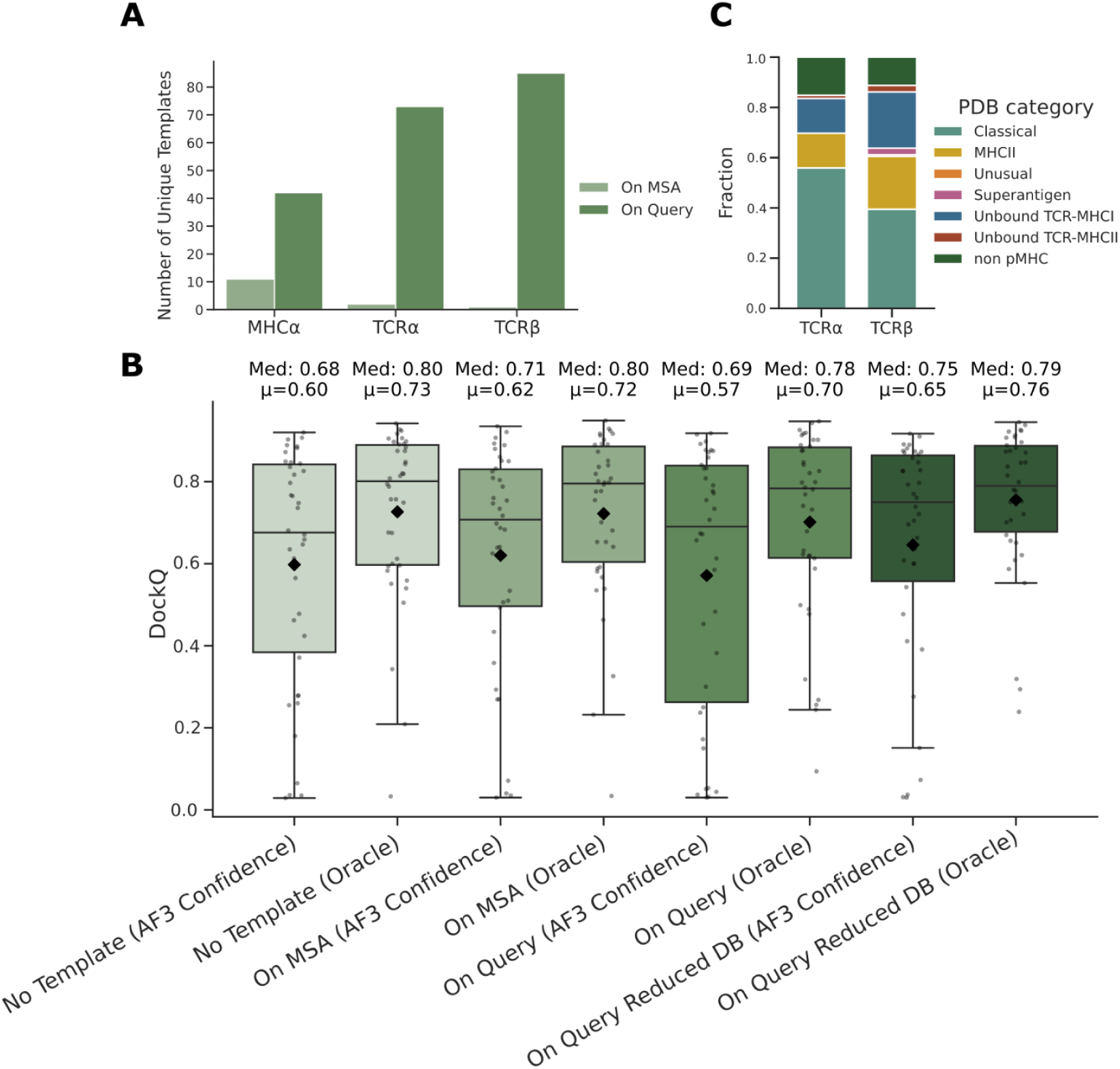
Effect of template search strategy and template database composition on template diversity and model quality. A) Number of unique templates selected per chain across the dataset, comparing template search performed on the full MSA profile versus the query sequence alone. B) Effect of template search strategy on model quality. Each datapoint has 50 models, generated from 10 seeds and 5 diffusion samples per seed, using unpaired MSA only and template search performed using either the full MSA profile or the individual chain sequence. The best model per datapoint was selected by max-pooling according to either DockQ (Oracle) or AF3 confidence from the model pool. C) Identity of the selected templates for TCRα and TCRβ, expressed as fractions across the benchmark dataset. Each point represents a TCR-pMHC complex, and the mean is shown by the diamond point.

Despite this increase in diversity, overall modeling accuracy decreased following this modification (see Figure 2B, On MSA vs On Query). We hypothesize that this decrease in accuracy is due to the template database composition, since the template-search on query leads to the selection of more diverse templates and therefore increases the reliance of the modelling on the quality and relevance of the retrieved templates. Consistent with this, approximately 30% for TCRa and 35% for TCRb of the selected templates were atypical TCR structures such as non-pMHC TCRs or unbound-TCRs, which do not recapitulate correctly the bound TCR-pMHC conformation and can therefore inform more poorly of the bound conformation (Figure 2C). Moreover, between 14% and 21% of the templates corresponded to TCR-MHC class II structures despite our system of interest being MHC class I-restricted. We tested whether restricting the template database to TCRs bound to MHC-I structures only would improve performance, but this did not change modeling accuracy (Supplementary Figure 1A), suggesting that templates of TCRs bound to MHC-II may still provide useful chain-level structural information. These results motivated the construction of a curated, TCR-pMHC specific template database, containing both MHC I and MHC II restricted structures but excluding all non-pMHC and unbound TCRs (see methods for details on database construction).

This resulted in a curated database consisting of 3,051 templates of MHC molecules and TCRs bound to pMHC (see Template database reduction in Methods for details). Combining this curated, TCR-pMHC specific template database with the query-based template search improved performance on our benchmark dataset relative to query-based search against the default, full AF3 template database (Figure 2B, On Query vs On Query Reduced DB). This improvement was concentrated in the tail of the Oracle DockQ distribution, reflecting a reduction in the number of low-quality predictions, with only 3 structures falling below the medium quality threshold (DockQ = 0.49) when using the curated database, compared to 7 structures below this threshold when using the full database (Supplementary Figure 1B).

The curated database also narrowed the gap between the Oracle-best model and the model selected by AF3’s own confidence metric, going from a median difference of 0.09 when using the full database to 0.04 when using the curated version, indicating that the improved template set produces models that are not only higher quality on average, but also easier to identify from confidence scores alone.

Together, these results suggest that while query-based template search is important for diversifying and tailoring template selection to each query, the composition of the template database itself is a significant driver of prediction quality, highlighting the importance of having a high quality template database. Throughout the remainder of this work, we will apply query-based search on the reduced database for template selection.

#### 1.3 The adapted pipeline successfully recapitulates TCR-pMHC complex geometry

To evaluate the structural quality of the models produced by our TCR-pMHC-specific AF3 pipeline, we compared models selected via Oracle or AF3 confidence against the reference structures across several structural metrics (see Methods for details), thereby quantifying the pipeline’s upper performance bound (Oracle) and the accuracy of AF3 confidence as a model-selection criterion.

Under Oracle selection, models showed high overall quality, with a median global TM-score of 0.98, and high per-chain lDDT, indicating reliable chain-specific modeling. TCR-peptide interface accuracy was lower, with the Oracle achieving a median iTM-score of 0.83. This indicates that the interface region is modeled with lower accuracy than the surrounding structure, although it remains high in absolute terms (see Figure 3A, Oracle). Under AF3 confidence based selection, both overall and chain-level quality remained high, with a median global TM-score of 0.97.

**Figure 3:**
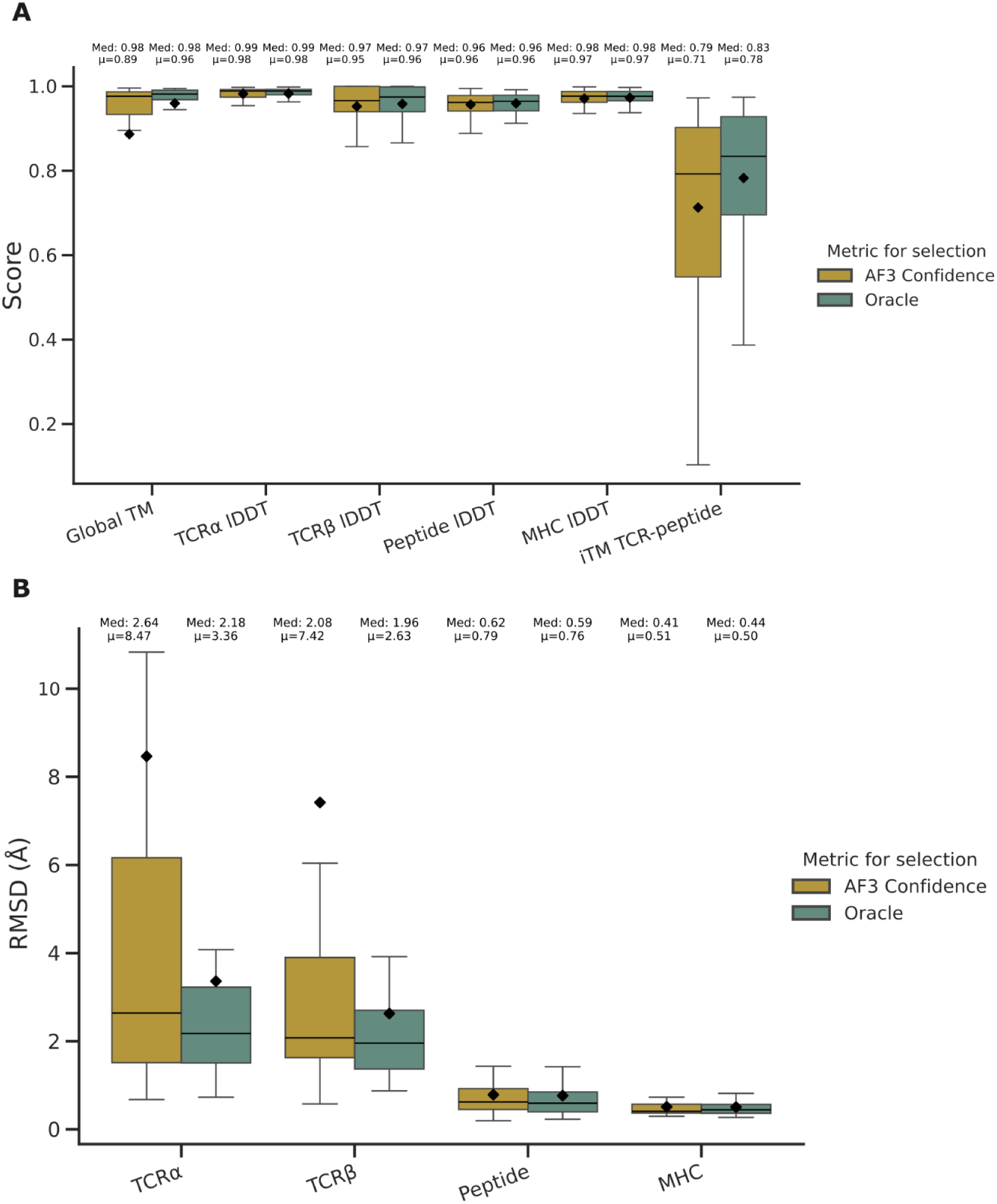
Structural metrics for the top-ranked model per datapoint (n = 38), comparing models selected using Oracle or AF3 confidence against the solved structure. A) Structural metrics assessing fold quality at the global level (global TM), chain-specific level (chain lDDT) and TCR-peptide interface level (iTM TCR-peptide). Higher values indicate better model quality. B) Structural metrics assessing docking accuracy relative to the MHC, shown as the RMSD of each chain when superimposing model and solved structures on the MHC. Lower values indicate more accurate docking. In both panels, the mean is indicated by the diamond point.

However, interface accuracy dropped substantially (median iTM score of 0.79), with AF3 confidence failing to consistently identify the best modeled interfaces (Figure 3A, AF3 Confidence).

We next examined TCR-pMHC docking geometry using the RMSD of each chain after MHC-based alignment (see Methods for details). This analysis showed that the peptide most often was correctly positioned within the MHC binding groove across models, under both Oracle and AF3 confidence selection strategies (Figure 3B, Peptide). Under Oracle selection, the TCRα and TCRβ chains were also confidently placed relative to the MHC, with mean/median RMSD values of 2.63/1.96 Å and 3.36/2.18 Å respectively. In contrast, under AF3 confidence based selection, TCR positioning errors were substantially higher, with mean/median RMSD values of 7.42/2.08 Å and 8.47/2.64 Å for TCRα and TCRβ respectively (Figure 3B, TCRα and TCRβ).

Visual inspection and quantification of a representative model at the TCR-pMHC interface confirmed this pattern with Oracle selection showing good overall backbone alignment with the reference structure, while the AF3 Confidence selected model showed significant backbone shifts (see Supplementary figure 2). Further, in 5% of the cases (2 of 28), AF3 confidence selected a model in which the TCR was not in contact with the peptide.

Overall, these results indicate that while the pipeline generates models with accurate chain-specific fold and TCR-pMHC docking geometry, AF3 confidence does not reliably identify these models. Therefore, relying on AF3 confidence for model selection risks selecting an incorrectly docked complex.

#### 1.4 Exploring confidence metrics for identifying best AF3 model

The results above motivate the evaluation of alternative metrics that better capture model quality, such that model selection more closely approximates Oracle performance.

Our finding that AF3 confidence selects a model without TCR-peptide contact in 5% of complexes, despite a correctly docked model being available in the pool, motivated an additional filtering step prior to model selection. Here, we applied an interface-based filter removing all models lacking an interface between the peptide and either the TCRα or TCRβ chain (for details refer to methods) .This filtering step corrected the two data points described above, shifting their selected models from mispositioned TCRs to docking geometries closely resembling the Oracle structure (Supplementary Figure 3).

Following this filtering step, we evaluated how model quality under Oracle selection compared to selection using several AF3 derived confidence metrics (see Methods for metric details). For each of the 38 benchmark structures, we computed the difference between the Oracle DockQ and the DockQ obtained using each candidate metric. Metric performance was then summarized as the median DockQ difference across all structures, with lower values indicating closer agreement with Oracle-based selection. For all metrics evaluated, this median difference remained below 0.1, indicating that every metric selects models with a quality close to that of the Oracle selection (Figure 4). Visualizing the per-datapoint DockQ differences confirmed this overall similarity across metrics, but also showed that lower-median metrics had fewer data points exceeding a difference of 0.1 (Supplementary Figure 4).

**Figure 4:**
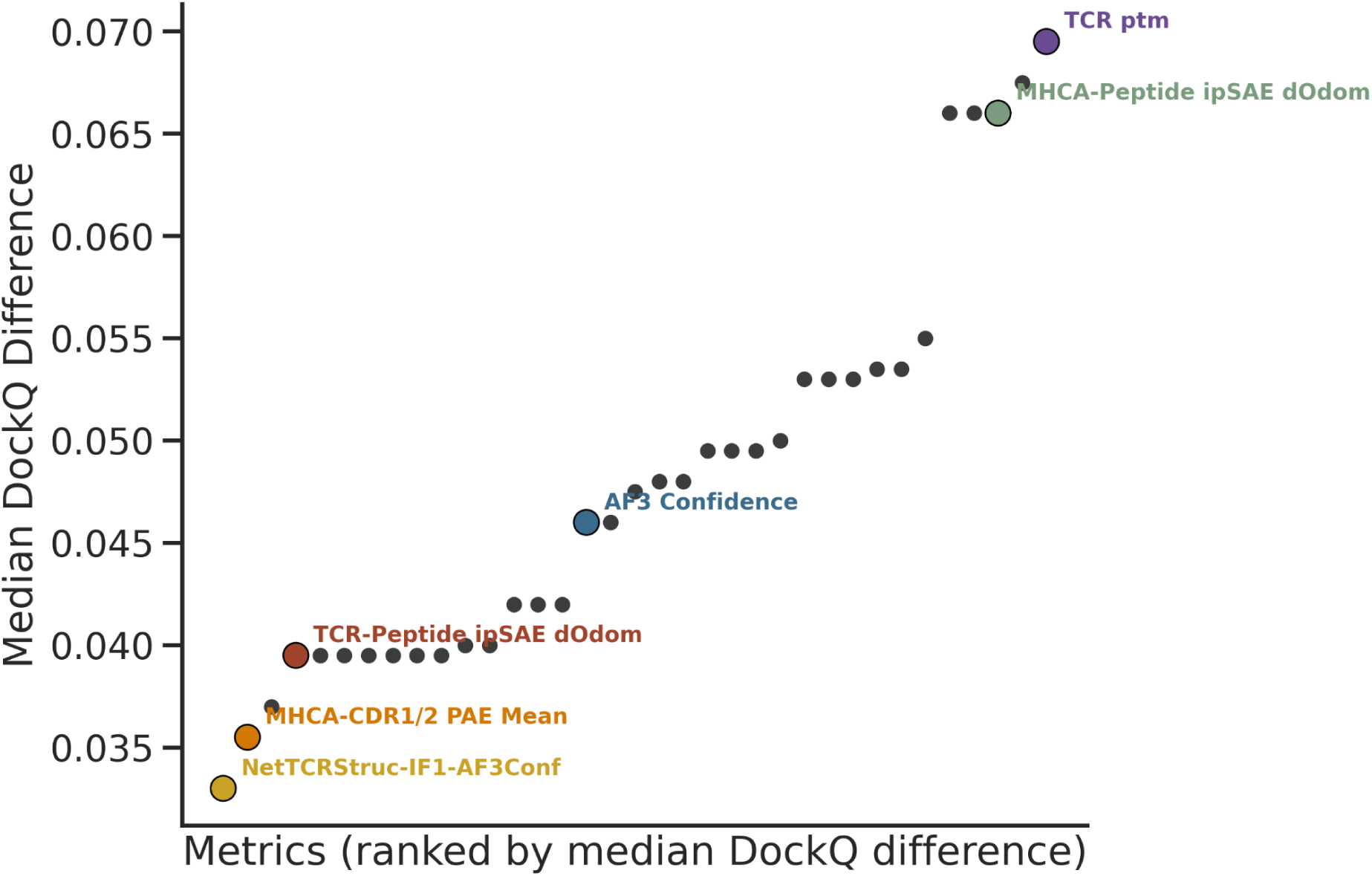
Median DockQ difference between Oracle selection and confidence metric based model selection. Confidence metrics are ranked from lowest to highest median DockQ difference relative to Oracle Selection. Four metrics are highlighted for subsequent analysis: NetTCRStruc-IF1-AF3Conf, MHCA-CDR1/2 PAE Mean, TCR-Peptide ipSAE d0dom and AF3 Confidence. Two of the worst-performing metrics, TCR pTM and MHC-peptide ipSAE d0dom, are also shown for comparison.

To identify patterns among metrics with either high or low agreement to the Oracle, we compared the 10 best-performing and 10 worst-performing metrics by median DockQ difference, out of the 35 metrics evaluated in Figure 4. Among the top 10 metrics, 90% (9 of 10) involved the TCR, either at the MHC-TCR interface (50%, 5 of 10) or the TCR-peptide interface (40%, 4 of 10), while only 1 of these TCR-related metrics appeared among the worst-performing. In contrast, the worst-performing metrics were dominated by those assessing the MHC-peptide interface (40%, 4 of 10) and those based on single-chain interactions (40%, 4 of 10). Global metrics were evenly split between the two groups, accounting for 10% of both the top and bottom 10, with the remaining global metrics falling in the middle range of the performance (Table 1).

**Table 1:**
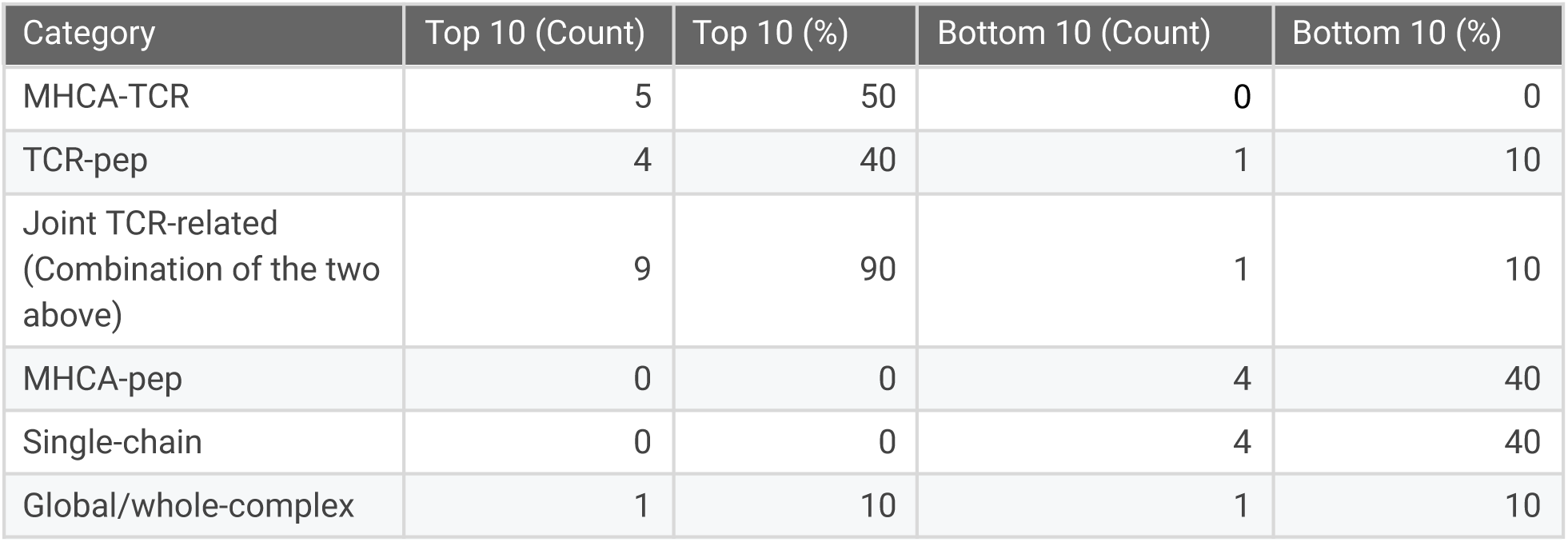
Composition of the 10 best-performing and 10 worst-performing metrics (ranked by median DockQ difference from Oracle, see Figure 4) are shown split by metric category. Results are reported as both counts and percentages.

Given this close agreement across metrics, we selected three top-performing metrics as model-selection metrics for downstream analyses: one derived from NetTCRStruc (NetTCRStruc-IF1-AF3Conf), one involving MHC-TCR interactions (MHCA-CDR1/2 PAE Mean), and one involving TCR-peptide interactions (TCR-Peptide ipSAE d0dom). We also included AF3 confidence in downstream comparisons, as a reference for the default model-selection approach (Figure 4).

#### 1.5 Identifying the best metric to stratify binders from non-binders

After establishing that our pipeline can generate and identify accurate TCR-pMHC structures, we next evaluated whether it could also stratify binders from non-binders. To do so, we constructed artificial negatives for the 38 structures of our benchmark by matching each pMHC with five TCRs known to bind other pMHCs (see methods), resulting in a dataset of 38 batches each containing one positive and 5 negative TCR-pMHC complexes. We then used the four metrics identified in the previous section to identify the best model for each complex (model selection).

Next, each selected model was scored using a series of metrics, and each metric evaluated based on its ability to distinguish true binders from negatives within each batch (target selection), under the hypothesis that only true binding complexes would resemble correct TCR-pMHC structures, and therefore receive higher scores from our selected confidence metric compared to the mis-matched negatives. We evaluated this using a metric called TPR (see materials and methods), which quantifies how many negatives are ranked above the true positive within each batch. Finally, the overall performance of a given metric is quantified as the mean TPR across batches (see Figure 5). The result of this analysis clearly demonstrated that TCR-peptide interaction metrics were the best performers for binder identification, achieving mean TPR values between 0.83 and 0.85 across the 38 batches. Metrics based on MHC-TCR interactions, although useful for model selection (see Figure 4 and Table 1), fell in the intermediate range for binder discrimination, with TPR values between 0.80 and 0.83. Per-chain confidence metrics performed worst overall, with TPR below 0.7 in all cases. Of note, NetTCRstruct derived metrics performed comparably to AF3 based metrics (See supplementary Figure 6), despite being trained on AF2.3 derived models rather than AF3.

**Figure 5:**
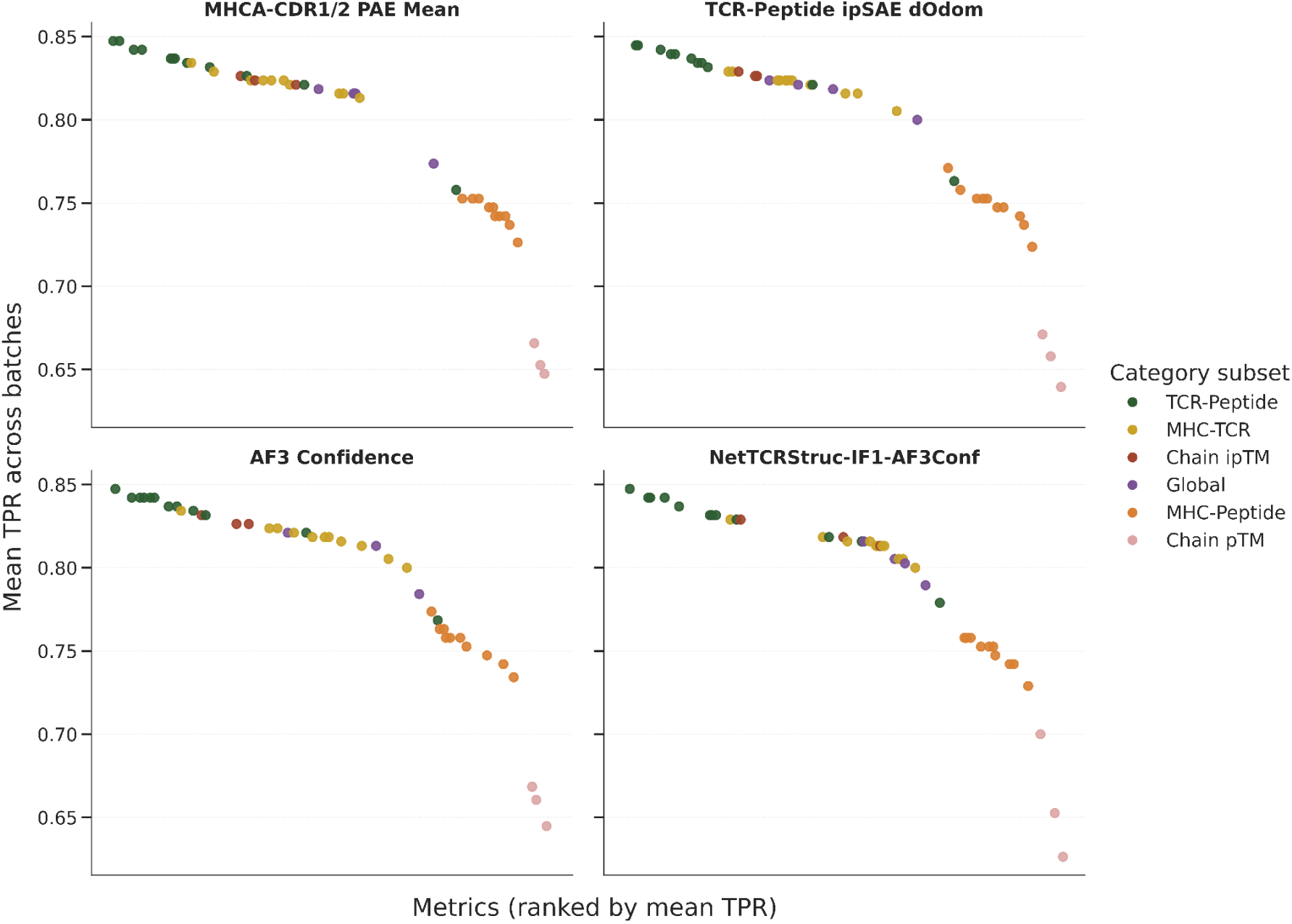
Performance of candidate metrics for target-selection. For each model-selection metric (facets), performance is evaluated as the mean TPR across the 38 batches, using each candidate metric for target selection. Metrics are ranked by mean TPR. Each of the four facets shows the same set of target-selection metrics, with each point representing one metric, colored by its category as indicated in the legend. TCR-peptide, MHC-TCR and MHC-peptide categories group metrics by the interface they capture. Chain ipTM and chain pTM categories group single-chain metrics (TCR, peptide or MHC ipTM or pTM). Global metrics (AF confidence, ipTM and pTM) consider the complex as a whole.

Together, these results indicate that TCR-peptide interface-based metrics can stratify binders from non-binders within our pipeline, with TCR-pep ipSAE d0dom being among the best-performing metrics. We therefore selected TCR-peptide ipSAE d0dom for target selection in all subsequent analyses, applying it in combination with two model selection strategies,TCR-peptide ipSAE d0dom itself, and the mean PAE between MHC and CDR1/2, respectively.

### Prediction of T Cell specificity

#### 2.1 The NetTCRfold pipeline outperforms state-of-the-art methods in TCR-pMHC specificity prediction

To further evaluate the pipeline’s ability to predict TCR-pMHC specificity, we assessed its performance on the Immrep2025 dataset (see Methods). As described above, the two metrics, the TCR-pep ipSAE d0dom, and the mean PAE between MHC and CDR1/2 residues, were used to select the best model per datapoint. Target selection was then inferred in both cases using the TCR-peptide ipSAE d0dom score as the target value.

Figure 6 shows the result of this analysis, demonstrating that our pipeline performed at par with the top-performing competition submission (AF3 Bradley Clustered), which relied on an additional clustering step to improve performance. Across the 18 private peptides, our pipeline achieved a median AUC0.1 of 0.61 under both model-selection strategies, with mean AUC0.1 values of 0.60 (MHC-CDR1/2 PAE mean for model selection) and 0.59 (TCR-peptide ipSAE d0dom for model selection). Our pipeline also outperformed Chai-1-based methods, as well as sequence-only approaches such as NetTCR.

**Figure 6:**
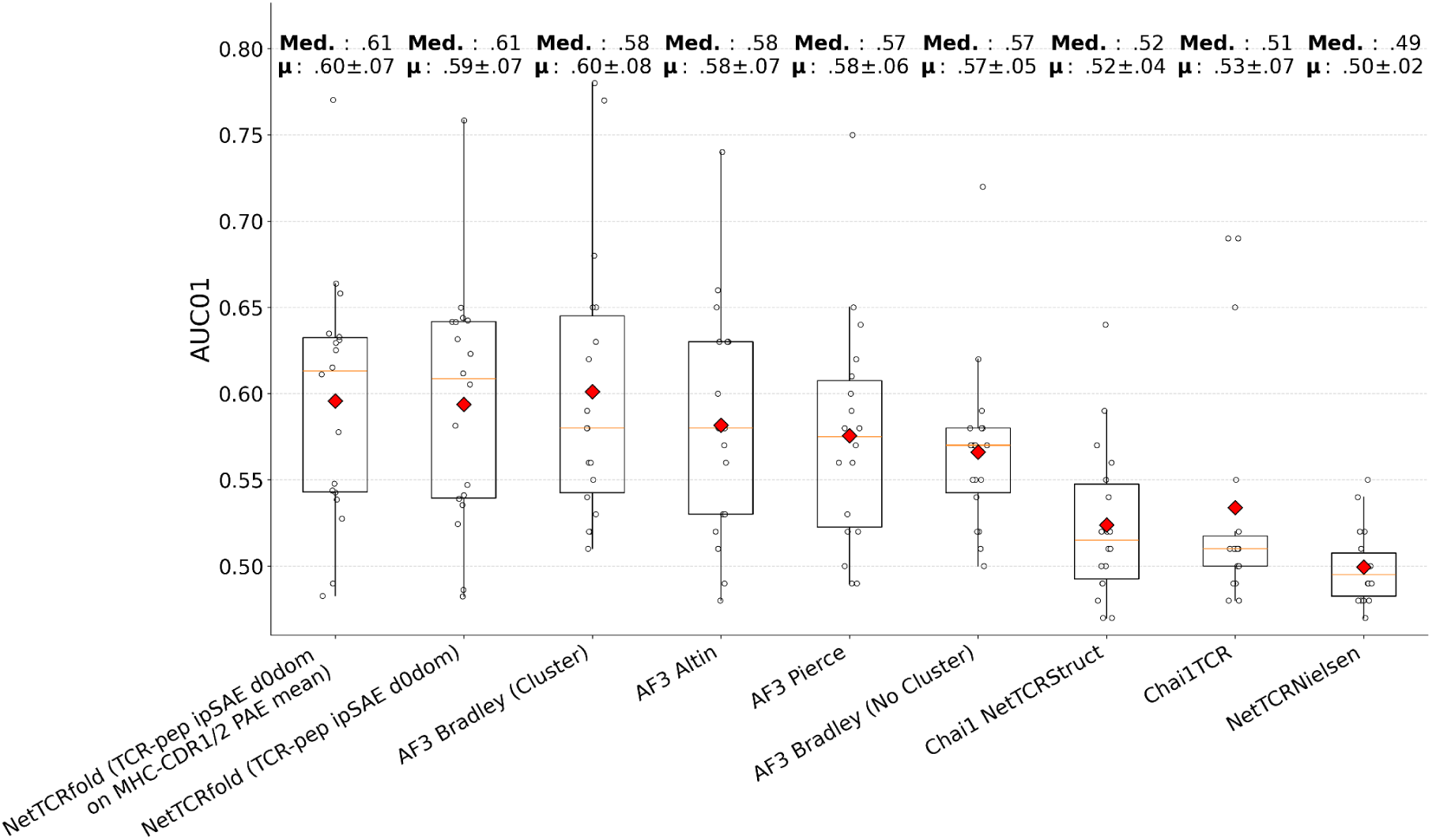
Per-peptide AUC0.1 across the 18 private peptides for our pipeline compared to top-performing Immrep2025 methods, including AF3-based methods (with and without clustering), Chai-1-based methods, and sequence-only. Dots show per-peptide AUC0.1 values and diamonds indicate the mean AUC0.1 per method.

At the HLA level, our method was the top performer for HLA-A*02:01. For HLA-B*40:01, performance was comparable to other AF3-based methods, but lower than TCRdock (AF3 Bradley) (Supplementary figure 5A). A full per-peptide breakdown of AUC0.1 across all private peptides and evaluated methods is provided in Supplementary figure 5B. Overall, our pipeline shows consistently strong performance across HLA-A*02:01 peptides, while performance across HLA-B*40:01 varies by peptide.

#### 2.2 NetTCRfold shows improved TCR-pMHC binding specificity prediction over AF2

Given these encouraging results, we continued the evaluation of TCR specificity prediction on the sequence dataset introduced by Deleuran et al. ^21^, which consists of true binders and artificial negatives generated by swapping TCR and pMHC pairs. This dataset consists of 2,945 binding complexes across 26 peptides, with 5 swapped negatives generated for each TCR-pMHC complex, resulting in a total of 2,945 batches, each consisting of 1 positive and 5 negatives (see Methods for details).

The modeling pipeline was applied to each datapoint. Model selection and target selection were then performed using four individual metric combinations: (1) TCR-peptide ipSAE d0dom for both model and target selection; (2) TCRα-peptide ipSAE d0dom for both model and target selection; (3) TCRβ-peptide ipSAE d0dom for both model and target selection; and (4) MHC-CDR1/2 PAE mean for model selection, combined with TCR-peptide ipSAE d0dom for target selection. Combinations restricted to a single TCR chain were included to examine each chain’s independent contribution to specificity prediction. For each of these four combinations, TPR performance metric was calculated per batch, the batches sorted by their highest target-selection score, and a cumulative TPR curve computed across batches (for details refer to Methods).

For comparison with AlphaFold 2.3, we evaluated the complexes modeled by Deleuran et al., using either AF2.3’s own confidence metric or the NetTCRstruct derived metric (GNN-IF1-AF) for model selection and target scoring. We additionally evaluated the AF3 confidence metric on our AF3 models, to assess its ability to distinguish binders from non-binders.

All four AF3-derived metrics outperformed both AF2.3-based scoring approaches and AF3’s own confidence metric (Figure 7A). Over the first 1,000 batches (34% of the dataset), AUC ranged between 0.84 and 0.88 for our metrics, compared to 0.71 for AF2.3 scored with its own confidence, 0.76 for AF2.3 scored with NetTCRstruct, and 0.75 for AF3 scored with its own confidence (Figure 7B). It is also worth noting that the AF3 confidence metric alone did not outperform AF2.3 scored with NetTCRstruct, highlighting the need for a metric tailored to TCR-peptide interactions when predicting specificity, rather than a general purpose confidence score.

**Figure 7:**
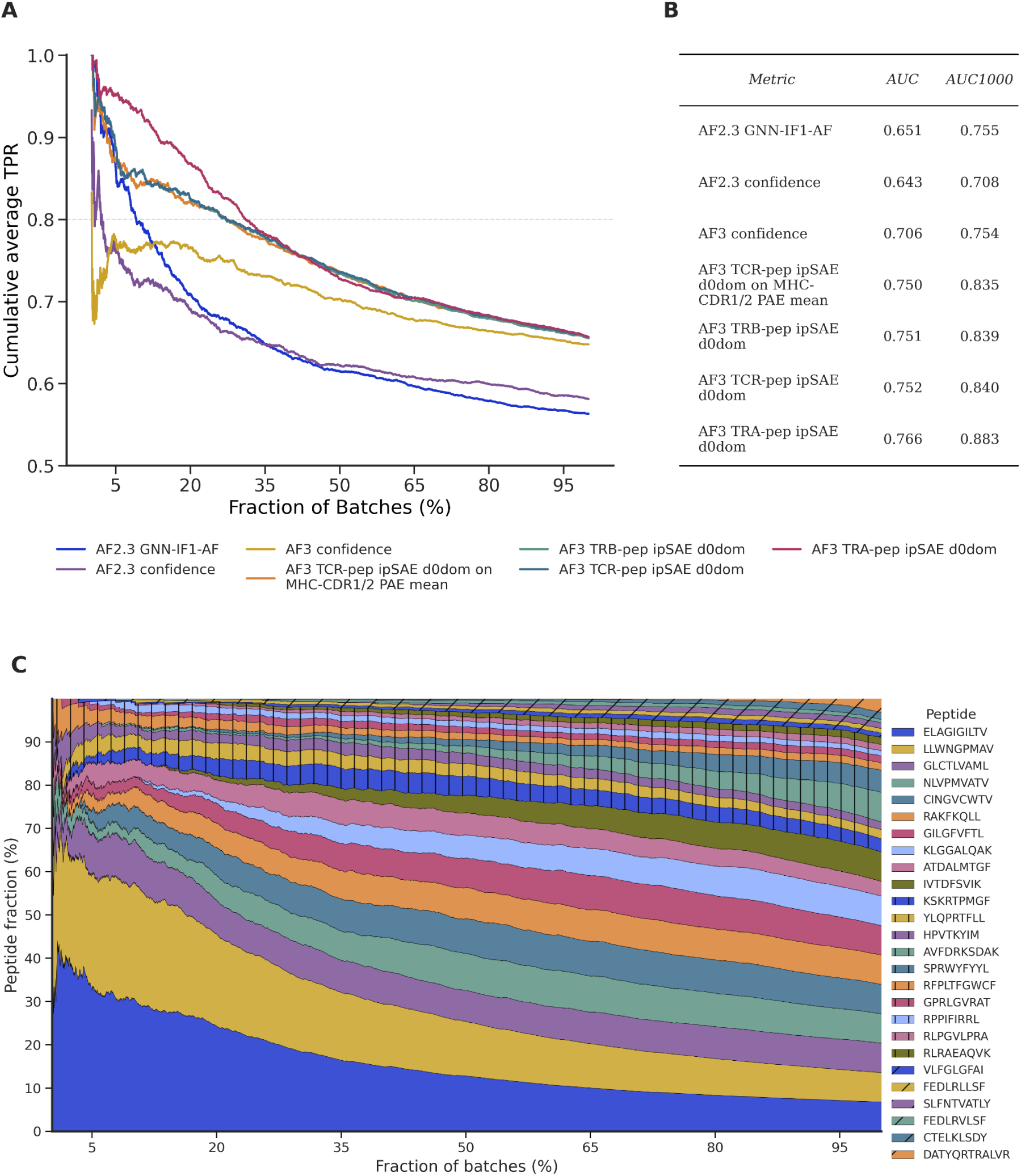
Specificity prediction performance across methods and metrics, and underlying peptide composition. A) Cumulative average TPR curve across batches, with batches sorted by maximal intra-batch score for each metric and displayed as a fraction of total batches. The legend indicates the metric used for target selection. If a different metric was used for model-selection, it is indicated by appending “on {metric to model selection}” to the metric name (see Methods for metric definitions). B) Table showing the AUC across all batches (AUC) and the AUC restricted to the first 1,000 batches (AUC1000), for every metric in panel A. C) Cumulative proportion of peptides as a function of batch fraction, with batches sorted by their maximal intra-batch TCRα-peptide ipSAE d0dom score.

Among our four metrics, the TCRα-only variant performed best over the first 1,000 batches, with an AUC of 0.88, compared to 0.84 for both TCRβ-only and the combined TCR score. This early advantage was not maintained across the full batch dataset, as all four metrics converged to an overall cumulative TPR of 0.66. To investigate this pattern, we computed a windowed cumulative TPR using a 500-batch rolling window rather than all preceding batches. This confirmed that TCRα-peptide outperforms the other metrics in top-scoring batches, but is surpassed by TCR-peptide and TCRβ-peptide metrics after roughly 20 to 25% of batches (Supplementary Figure 8A). Examining the distribution of peptides among these top 20% batches ranked by their TCRα-peptide ipSAE d0dom score, we found that more than 50% were associated with the ELAGIGILTV and LLWNGPMAV peptides (Figure 7C). This observation is consistent with previous reports showing a prevalence of the TRAV12-2 gene for TCRs specific to these two peptides, with CDR1α playing a crucial role in peptide recognition ^43,44^.

Overall, our pipeline achieved a cumulative TPR above 0.8 for 35% of the complexes, corresponding to an average rank within top-2 within each batch. In comparison, only 5% of complexes reached this threshold when ranked by AF2.3 confidence, and around 15% when ranked by NetTCRstruct. Together, these results align with earlier findings and suggest that the proposed pipeline can separate true binders from swapped TCR-pMHC pairs in cases of high structural quality, provided the scoring metric is tailored to capture TCR-peptide interactions^21^.

### Effect of noise on TCR-pMHC binding prediction

The results in Figure 7 above demonstrate that despite the improvements achieved by the refined modeling pipeline, a tail of predictions still show limited performance with TPR values falling below 0.8. Previous studies have suggested that a substantial proportion of the publicly available TCR-pMHC data consists of noise, and we hypothesize that this could be a contributing factor to the reduced binding prediction performance observed in some batches ^19^. If a significant fraction of the positive data is in fact a noisy, non-binding label, the corresponding batch would effectively consist of 6 non-binders, leaving no true signal to identify and resulting in a near random TPR.

#### 3.1 Label noise limits TCR-pMHC specificity prediction performance

To assess to what degree noise was driving the reduced prediction quality, we evaluated TCR-pMHC binding prediction on a subset of the dataset published by Messemaker et al. ^19^, which provides in-vitro denoised labels for two peptides. To do so, we subset the dataset introduced in the previous section to only include the batches where the positives were investigated in the Messemaker study, resulting in 96 batches: 54 for YLQPRTFLL (35 where the positive was confirmed as true binder and 19 where the positive was as noise (false positives)) and 42 for GLCTLVAML (28 where the positive was confirmed as true binder and 14 as false positives) (see Methods for further detail on dataset construction).

We next hypothesized that predictions on the denoised dataset (batches with confirmed binders) would show improved TCR-pMHC binding identification, while the noisy data (batches with false positive binders) would show near random performance.

We first predicted binding across the entire subset without distinguishing true binders from noisy labels, emulating the scenario present in our global dataset (Figure 8, full). Overall, predictive accuracy for these two peptides was higher than observed across the full dataset, with most batches above a cumulative TPR of 0.8 and AUCs of 0.89 and 0.90 for our selected metrics (see table 2, full dataset). We then split the dataset into noisy TCR-pMHC complexes (n = 33) and true binders (n = 63) (Figure 8, denoised and noise). After denoising, binding was predicted with a cumulative TPR above 0.9 across all batches, with an overall AUC of approximately 0.95 (see Table 2, denoised). For peptides labeled as noise, cumulative TPR remained below 0.8 across all batches, indicating limited difference in the signal between positives and negatives in those complexes.

**Figure 8:**
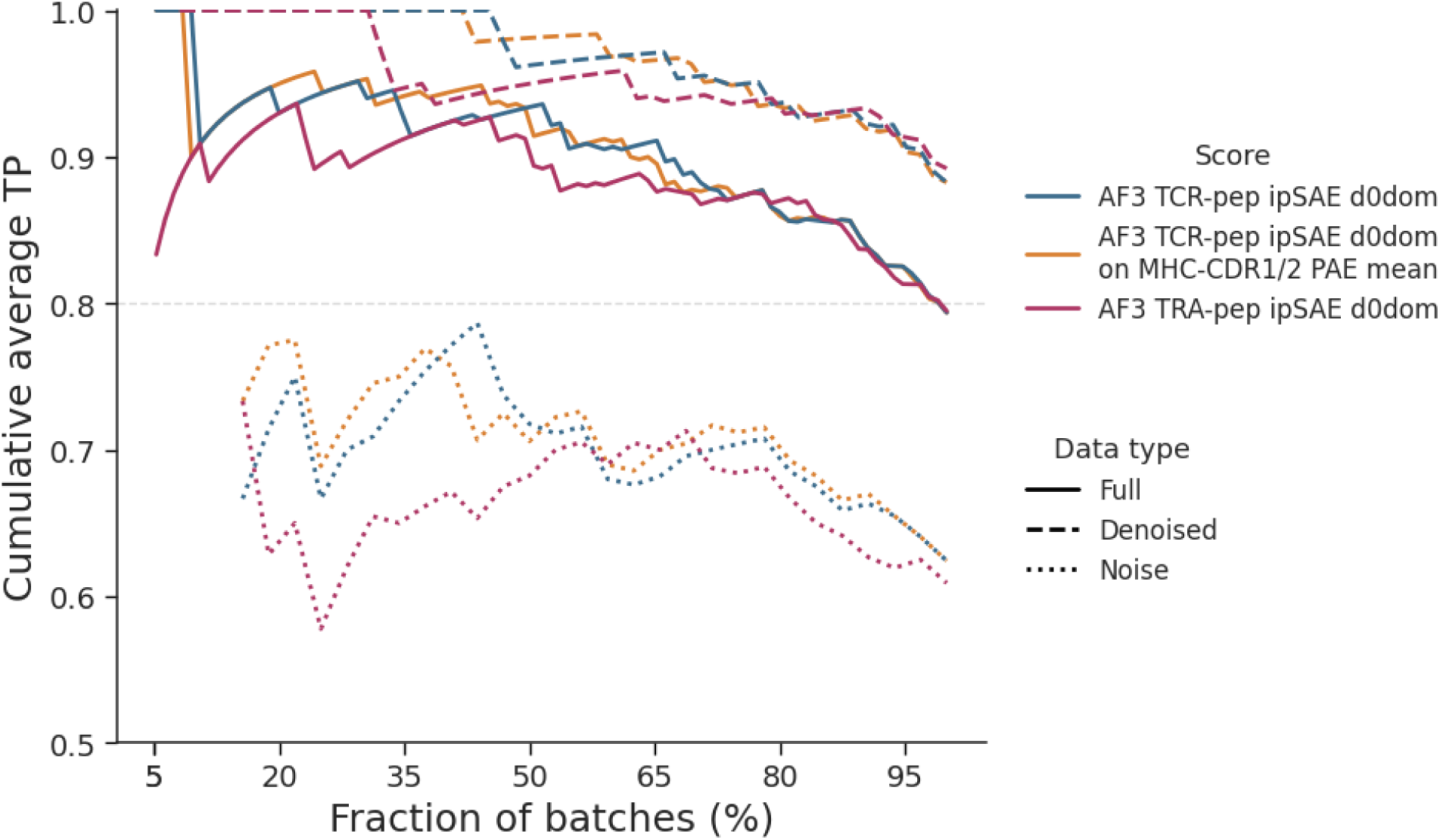
Effect of label noise on TCR-pMHC specificity prediction, evaluated on a subset of in-vitro denoised datapoints. Cumulative average TPR curve across batches, with batches sorted by maximal intra-batch score for each metric and displayed as a fraction of total batches. Line type indicates the full, denoised and noise-only datasets (see data type legend). Metric for target-selection is indicated in the score legend. If a different metric was used for model-selection, it is indicated by appending “on {metric to model selection}” to the metric name (see Methods for metric definitions).

**Table 2:** AUC of the cumulative TPR curve for the subset of the sequence dataset validated in the Messemaker study, by dataset subset (Dataset) and scoring metric (Score).

| Score | Dataset | AUC |
| --- | --- | --- |
| AF3 TRA-pep ipSAE d0dom | full | 0.874 |
| AF3 TCR-pep ipSAE d0dom | full | 0.895 |
| AF3 TCR-pep ipSAE d0dom on MHCA-CDR1/2 PAE mean | full | 0.896 |
| AF3 TRA-pep ipSAE d0dom | denoised | 0.938 |
| AF3 TCR-pep ipSAE d0dom | denoised | 0.951 |
| AF3 TCR-pep ipSAE d0dom on MHCA-CDR1/2 PAE mean | denoised | 0.952 |
| AF3 TRA-pep ipSAE d0dom | noise | 0.639 |
| AF3 TCR-pep ipSAE d0dom | noise | 0.676 |
| AF3 TCR-pep ipSAE d0dom on MHCA-CDR1/2 PAE mean | noise | 0.685 |

These results suggest that once the dataset is denoised and only true binders remain, the pipeline can accurately distinguish binders from non-binders. This result however also underlines that noise is a major determinant of predictive performance.

#### 3.2 Denoising TCR-pMHC specificity labels substantially improves predictive accuracy

After observing the influence of label noise in the small batch subset derived from the Messemaker study, we expanded the analysis to the full Deleuran et al. sequence dataset. Here, we ran a sequence-based denoising clustering algorithm, termed TCRdenoise, to identify potential wrongly labeled positive data points. In short, the denoising method takes as input a matrix of pairwise TCR distances calculated from a given sequence similarity metric, and performs hierarchical clustering optimizing a Silhouette score adapted to handle singletons and outliers. From the optimal clustering solution, non-clustered TCRs are defined as noise. An example of such a clustering solution is illustrated for IVTDFSVIK (IVT) in Figure 9A (for details about the clustering method refer to Methods and supplementary Figure 7). This denoising step classified 1,295 (44%) of the 2,945 datapoints in the full dataset as noise and the remaining 1,650 as binders.

**Figure 9.**
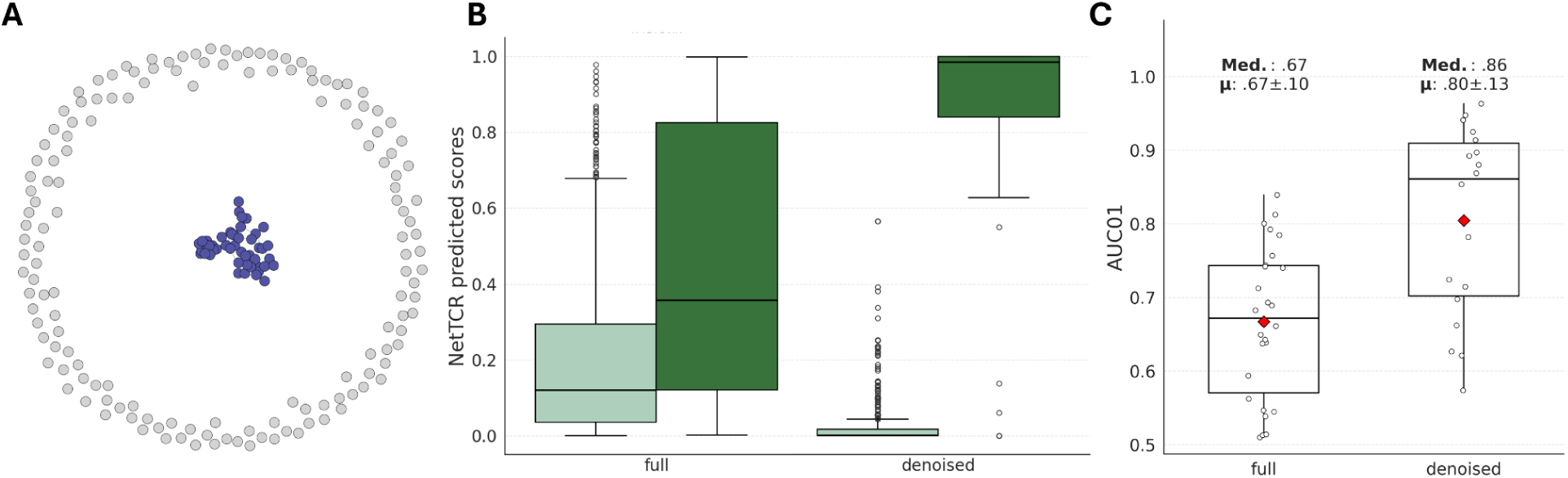
A sequence-based denoising approach A) Clustering solution for IVTDFSVIK, datapoints coloured grey are TCRs deemed noise by the denoising-algorithm and the datapoints colored blue are defined as binding TCRs. B) NetTCR prediction score distribution from models trained and evaluated on either a full dataset and on a denoised dataset. Light green boxes are the swapped negative predictions score distribution and the dark green are the positive datapoints distribution C) Distribution of the two models AUC01 scores for each peptide.

As a first evaluation of the clustering results, we trained and evaluated NetTCR on two different datasets, the full dataset and the denoised dataset obtained from the clustering-based denoising algorithm described above. Of note, the denoised dataset was further reduced slightly to 1,496 datapoints excluding data for 8 peptides with less than 5 positive TCR per cross-validation partition (for details on the NetTCR model training, and data set preparation, refer to Methods). Figure 9B displays the distribution of NetTCR scores for positive and negative data points for the peptide IVT for the two models. This plot clearly shows that the model trained on the denoised dataset has more well separated prediction score distributions for the two classes compared to the model trained on the full dataset. Repeating this analysis across the full set of peptides leads to an improved score distribution separation across most peptides, resulting in an overall large increase in model predictive performance (see Figure 9C).

We then continued to the batch rank evaluation, and split the full sequence dataset into batches whose positive example was classified as a binder by the denoising algorithm (denoised dataset), and batches whose positive example was classified as noise (noise dataset), and generated the associated cumulative TPR curves.

Figure 10 shows the result of this analysis. Here, the denoised dataset showed an improvement in prediction quality, maintaining a cumulative TPR above 0.8 for the top 60% of the batches, compared to the 35% in the full dataset, an increase of more than 70%. In contrast, for the subset of batches classified as noise, prediction quality dropped below 0.8 TPR in the first 5% of batches, consistent with the pattern observed in previous sections. The tail of low quality predictions observed in the full dataset was further absent in the denoised dataset, further indicating that label noise was a driver of these low quality predictions. This improvement was also reflected in the AUC, which increased from around 0.75 in the full dataset to between 0.82 and 0.84 in the denoised dataset, an increase of ∼10% (see Table 3,). These results indicate that the presence of noise in the original dataset led to an underestimation of the pipeline’s true performance and that once label noise is removed, our pipeline identifies true binders with high accuracy.

**Figure 10:**
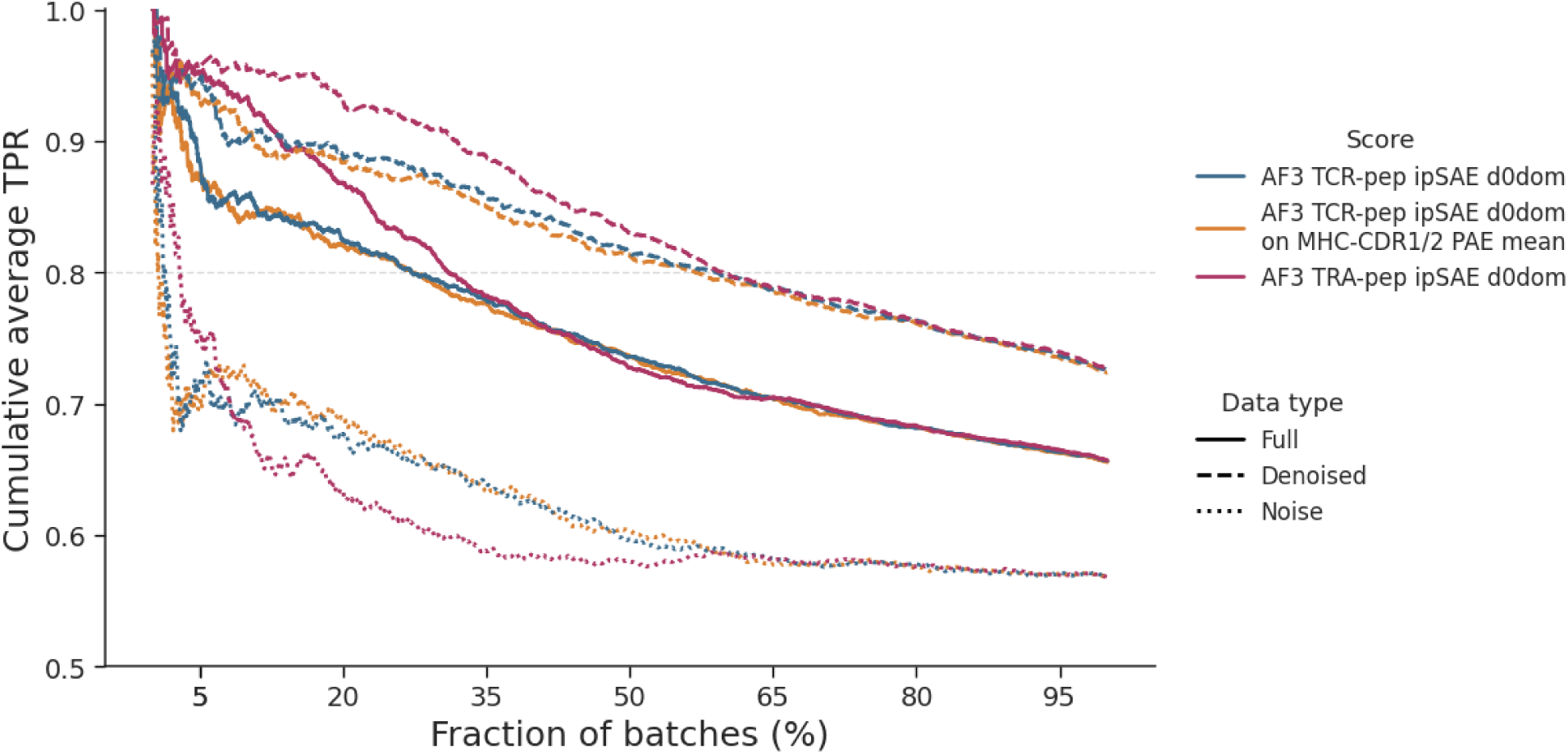
Effect of label noise on TCR-pMHC specificity prediction, evaluated on the full sequence database and denoised using a sequence based clustering algorithm. Cumulative average TPR curve across batches, with batches sorted by maximal intra-batch score for each metric and displayed as a fraction of total batches. Line type indicates the full, denoised and noise-only datasets (see data type legend). Metric for target-selection is indicated in the score legend. If a different metric was used for model-selection, it is indicated by appending “on {metric to model selection}” to the metric name (see Methods for metric definitions).

**Table 3:**
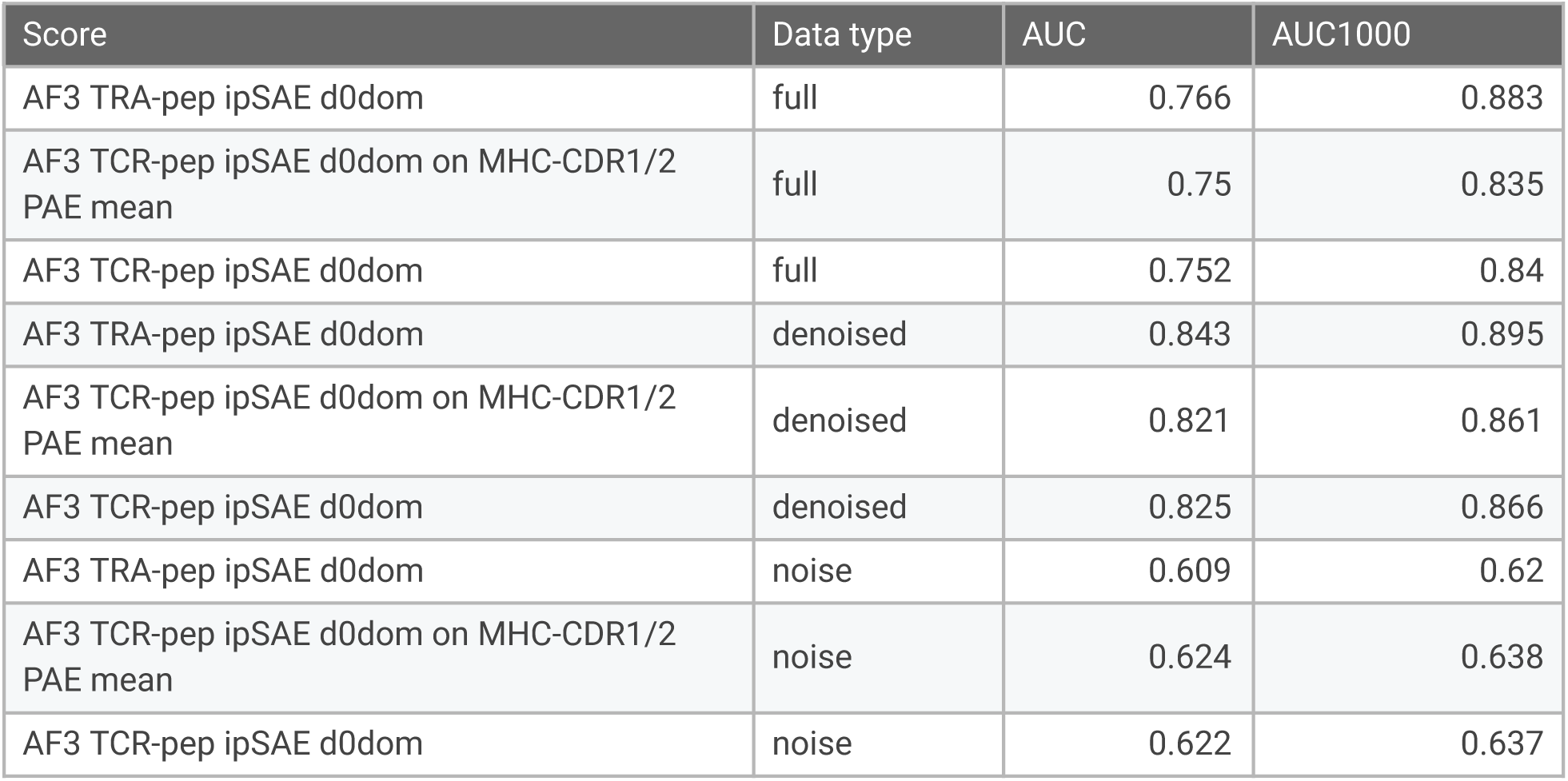
AUC of the cumulative TPR curve corresponding to the denoised sequence dataset, by dataset subset (Dataset) and scoring metric (Score).

| Score | Data type | AUC | AUC1000 |
| --- | --- | --- | --- |
| AF3 TRA-pep ipSAE d0dom | full | 0.766 | 0.883 |
| AF3 TCR-pep ipSAE d0dom on MHC-CDR1/2 PAE mean | full | 0.75 | 0.835 |
| AF3 TCR-pep ipSAE d0dom | full | 0.752 | 0.84 |
| AF3 TRA-pep ipSAE d0dom | denoised | 0.843 | 0.895 |
| AF3 TCR-pep ipSAE d0dom on MHC-CDR1/2 PAE mean | denoised | 0.821 | 0.861 |
| AF3 TCR-pep ipSAE d0dom | denoised | 0.825 | 0.866 |
| AF3 TRA-pep ipSAE d0dom | noise | 0.609 | 0.62 |
| AF3 TCR-pep ipSAE d0dom on MHC-CDR1/2 PAE mean | noise | 0.624 | 0.638 |
| AF3 TCR-pep ipSAE d0dom | noise | 0.622 | 0.637 |

## Code and data availability

The code and databases for the NetTCRfold pipeline can be accessed under: https://github.com/mnielLab/NetTCRfold-1.0.

## Discussion

In this work, we have introduced a refined AlphaFold3 TCR-pMHC class I structural modeling pipeline, termed NetTCRfold, and evaluated its ability to predict TCR-pMHC structures and binding specificity.

We first adapted the AF3 pipeline to incorporate TCR-pMHC domain-specific knowledge, under the assumption that this would improve modeling quality. This included modifications previously introduced by Yin et al.^11^, such as a domain-specific MSA database and query based template search, as well as two novel modifications: unpaired-only MSA alignment, and a curated TCR/MHC template database.

The main advantage of using domain-specific MSA databases was a large reduction in the computation time for structure generating, decreasing the time 54-fold (from 54 minutes per complex to 1 minute). Unpaired-only MSA outperformed AF3’s default setting, consistent with the fact that TCR chain pairing is not a product of co-evolution, so species-based cross-chain pairing does not reflect the system’s underlying biology. Restricting template search to the query sequence increased template diversity, but only translated into improved model quality when introducing a curated TCR/MHC template database, since the more diverse template search on its own resulted in the selection of TCRs bound to non-pMHC complexes or unbound structures.

Next, we found that AF3’s default confidence metric is not the best choice for either model selection or target selection. AF3’s default confidence metric often failed to identify correctly and optimally docked models from the pool of generated poses (model selection), even when such high quality models were present in the pool of poses. Here, we found that TCR-related metrics (either TCR-peptide or TCR-MHC) performed better. Also, for target-selection (i.e. selecting the best target for a given TCR), metric choice highly influenced the discrimination ability of the pipeline. We saw that using TCR-peptide interface metrics consistently improved specificity prediction, regardless of the metric used for model-selection. Combined, these results underline the critical importance of tailoring selection metrics to the task at hand, rather than relying on a general-purpose confidence score. Throughout this work, we found TCR-peptide ipSAE d0dom to be the best metric for specificity prediction, and either TCR-peptide ipSAE d0dom itself or the mean PAE between MHC and CDR1/2 to be the best for model selection. Across the different specificity datasets, both strategies gave highly similar performance, and for simplicity we therefore recommend using TCR-peptide ipSAE d0dom for both model selection and target selection.

We evaluated our pipeline against state-of-the-art TCR specificity prediction methods on the Immrep2025 dataset ^8^, and on a larger dataset introduced by Deleuran et al. ^21^. On Immrep2025, our pipeline performed at par with the top competition submission without requiring an additional clustering step. Performance was strongest for HLA-A*02:01, while performance for HLA-B*40:01 varied by peptide, consistent with this allele’s underrepresentation in public structure databases. On the Deleuran dataset, our pipeline outperformed AF2.3-based scoring approaches, including both AF2.3’s confidence metric and the refined NetTCRstruc score. Likewise, AF3’s default confidence metric was found to have sub-optimal performance, and the strongest performance was found when using a TCR-peptide interface metric for target selection. The overall conclusion from these benchmarks was, in line with earlier work ^21^, that TCR specificity prediction is feasible in the subset of cases where highly accurate structural models of the TCR-pMHC complex can be generated, and that this subset of cases can be greatly enlarged with the use of AF3 and refined model confidence metrics.

Investigating at the contribution of confidence scores of individual TCR chains in specificity prediction in the Deleuran dataset, a TCRα-peptide interaction metric was found to outperform both the combined TCR score and a TCRβ-peptide metric for two peptides (ELAGIGILTV and LLWNGPMAV). TCRs binding to these peptides are known to share a high preference for the TRAV12-2 gene, with the CDR1α playing a major role in peptide recognition through direct contact with the peptide N-terminus ^43,44^. This suggests that tailored metrics can be of value if some prior knowledge exists related to TCR-pMHC binding mode. However, in practice this chain-specific prevalence is not usually known, so we consider a combined TCRα and β metric the more broadly useful choice for specificity prediction.

Label noise also had a clear effect on specificity prediction. Public TCR-pMHC data are known to contain a substantial proportion of false-positive labels and we show that this noise leads to a substantial underestimation of true model performance for specificity prediction ^19^. This was shown by the improvement in predictive performance after removing noisy data points using first the small data set of Messemaker et al. with experimentally annotated noise labels, and next our sequence-based dataset, combined with a novel sequence-based denoising algorithm. In both cases, overall performance was increased by more than 10%, and the proportion of TCRs with accurate specificity prediction increased by more than 70%, when evaluated on the denoised data compared to the full dataset. Moreover was the performance on the noise data set consistently found to be close to random. These results demonstrate that the performance of structure-based TCR-pMHC specificity prediction most often is significantly higher than what has been reported in the literature, and we consider label noise an essential challenge to address, since the high noise level in current public data limits model-developement within both the structure and sequence domains.

It is critical to underline that our batch design, using one positive and five negatives, does not fully reflect real clinical settings, where a binding prediction study will typically involve a much larger set of TCR-pMHC candidates. Increasing the number of negatives would better reflect this setting, but due to computational and data constraints, we did not evaluate performance under this scenario here. We however do not expect this limitation to impact any of the general conclusions put forward in the work. Moreover, although predictive performance has increased relative to sequence based approaches, from close to random to some discriminative power (AUC=0.61), our findings suggest that these reported values may underestimate the model’s capabilities, since they do not account for label noise. Therefore, re-evaluating current methods on denoised data would better reflect how close structural approaches are to clinical applicability. Finally, while this work addressed the computational bottleneck of the MSA generation step, reducing runtime from 54 minutes per complex to around 1 minute, the inference step of these structural modeling approaches remains computationally expensive, taking approximately 10 minutes to generate 50 models per datapoint. Large scale structure generation therefore requires significant time and GPU resources, which limits the scalability of these methods to the large datasets required in clinical settings.

We see two main directions for future work. First, the label noise problem highlighted in this work motivates development of denoising strategies, combining in-vitro experimental validation with sequence and structure based approaches, such as the clustering method introduced here. The second direction concerns the overall modeling quality of TCR-pMHC complexes, as the ability to predict TCR specificity, as mentioned above, is fully contingent on this. Such improved model accuracy could potentially be improved by including contact restraints in the modeling framework ^45,46^, a feature not currently supported by AF3, or through domain-specific modifications such as TCR-pMHC model fine-tuning or joint pMHC template modeling, both earlier explored in AF2.3-based tools such as TCRdock ^9^ and TCRmodel2 ^11^.

In conclusion, we have introduced a modified AlphaFold3 pipeline tailored to TCR-pMHC structural modeling and demonstrated that it can be used to generate accurate models of TCR-pMHC complexes and predict TCR-pMHC binding specificity. The pipeline outperforms both AF2.3-based methods and other state-of-the-art AF3-based methods when used together with a target-selection metric focused on the TCR-peptide interface. Our results further show that label noise in public datasets currently limits the accurate evaluation of model performance, with specificity prediction improving substantially when denoised datasets are used.

## Funding / Acknowledgement

Research reported in this publication was supported by the Novo Nordisk Foundation Data Science Collaborative Research program under the grant NNF24OC0089619. The authors further acknowledge the funding from the Novo Nordisk Foundation (Grant ID: NNF25OC0105113) to use the Gefion AI Supercomputer, operated by the Danish Centre for AI Innovation (DCAI), for this work.

---

**Supplementary Figure 1:**
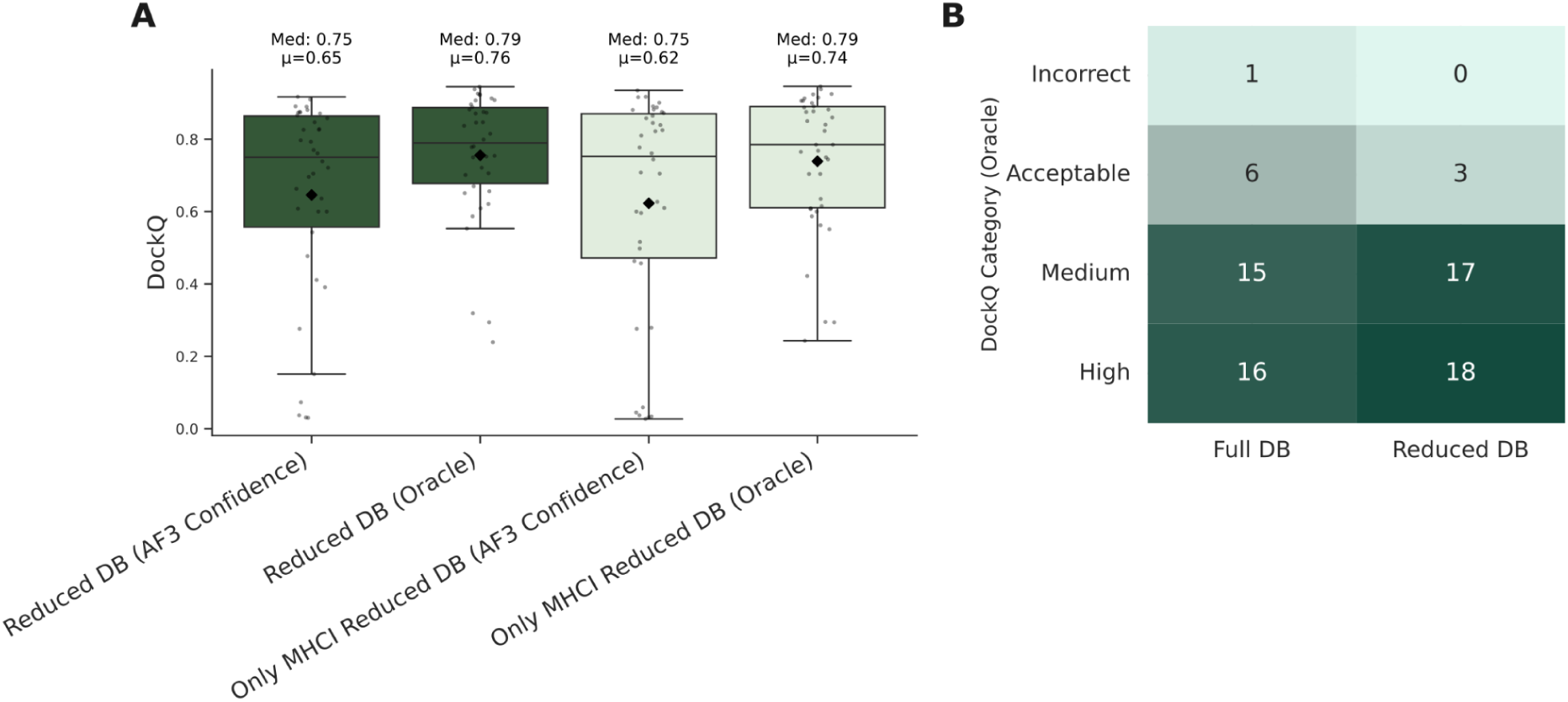
A) Effect of reducing the template database to MHC I related TCR complexes and MHC molecules. Reduced DB contains the reduced template dataset including also TCRs bound to MHC class II and MHC class II templates, while Only MHC I reduced DB removes these entries, and contains only MHC I related entries. B) Oracle DockQ shown by DockQ category when using the full template database or the curated database.

**Supplementary figure 2:**
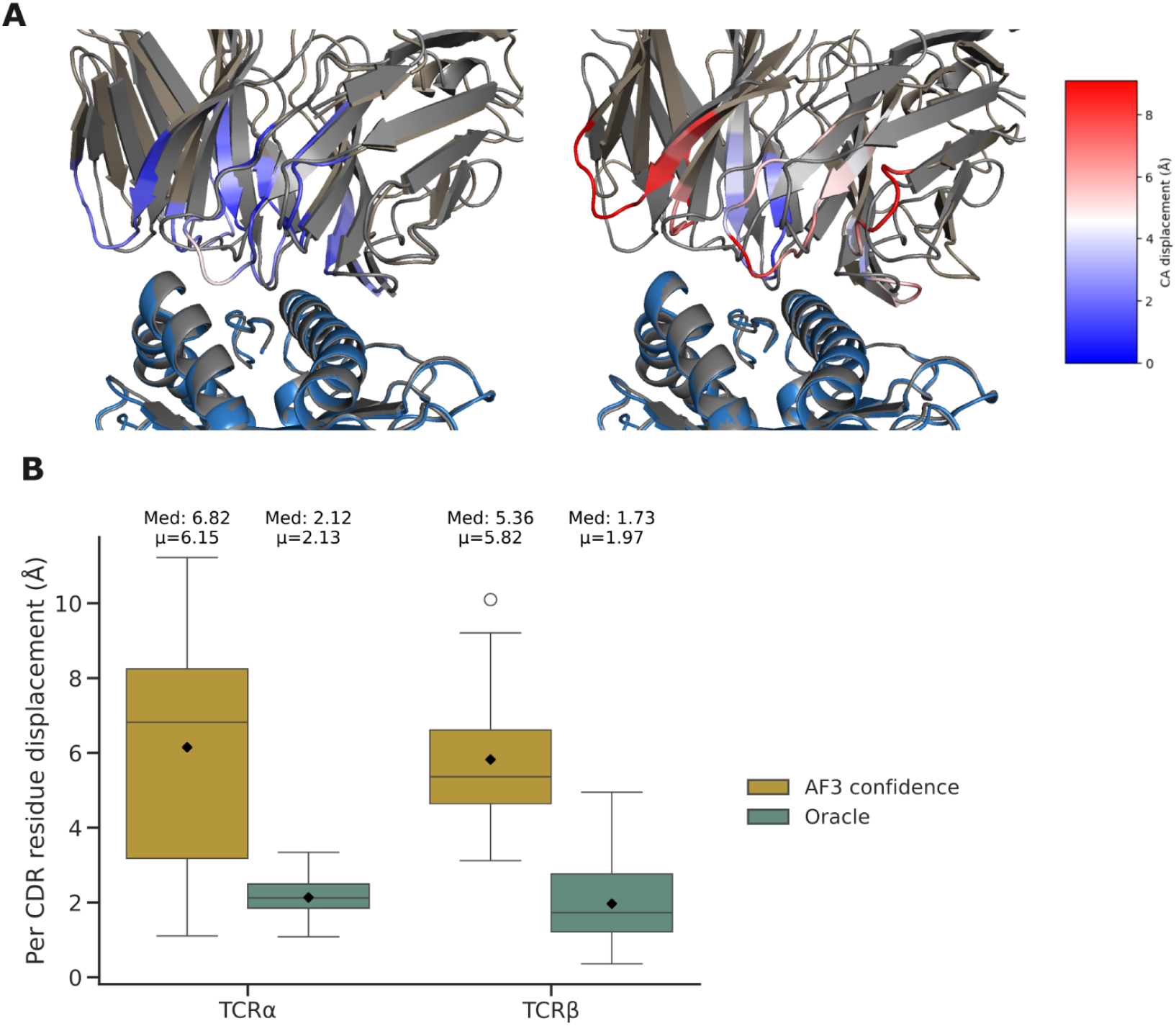
A) Per-residue displacement between model and solved structure at the six CDR loops, shown for the same datapoint (7rm4), comparing the model selected using Oracle and the model selected using AF3 confidence. Displacement was computed per Cα atom (see Methods) and restricted to the CDR loops. The color scale is shared between both structures for direct comparison. Solved structure is shown in gray and the model residues for which displacement was not obtained are shown in brown. Left: Oracle selected model, with aligned RMSD of 2.41Å and 3.1Å for TCRα and TCRβ respectively, close to the mean aligned RMSD under Oracle selection. Right: AF3 confidence selected model, with aligned RMSD of 7.42 Å and 8.47 Å for TCRα and TCRβ respectively, close to the mean aligned RMSD under AF3 confidence based selection. B) Per-residue CDR displacement for TCRα and TCRβ, shown separately for each selection metric, for the two representative models introduced in panel A.

**Supplementary Figure 3:**
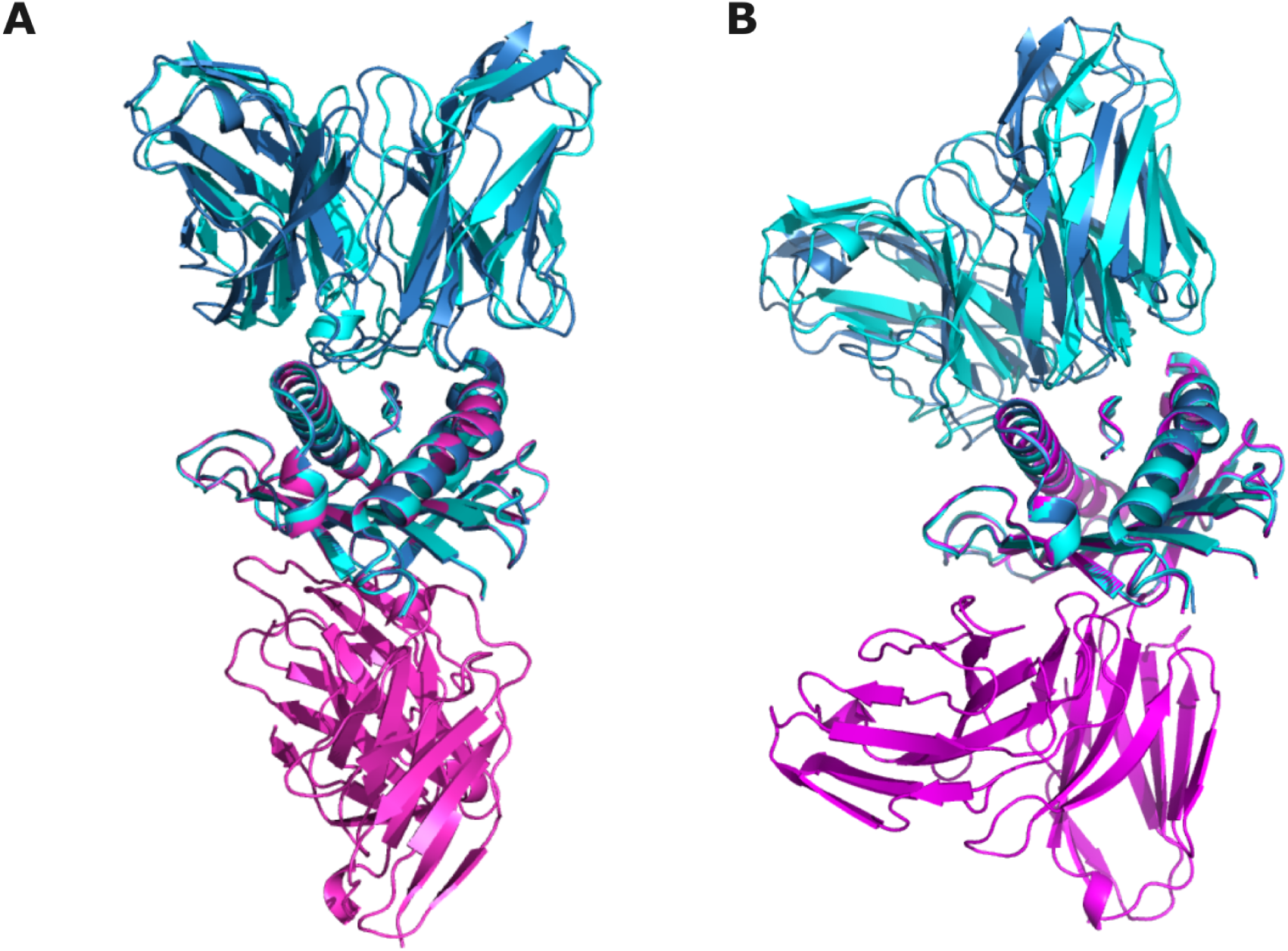
Models selected by Oracle (darkblue) and AF3 confidence before interface-based filtering (magenta) and after interface-based filtering (cyan) for 7rk7 (left) and 8ryo (right).

**Supplementary figure 4:**
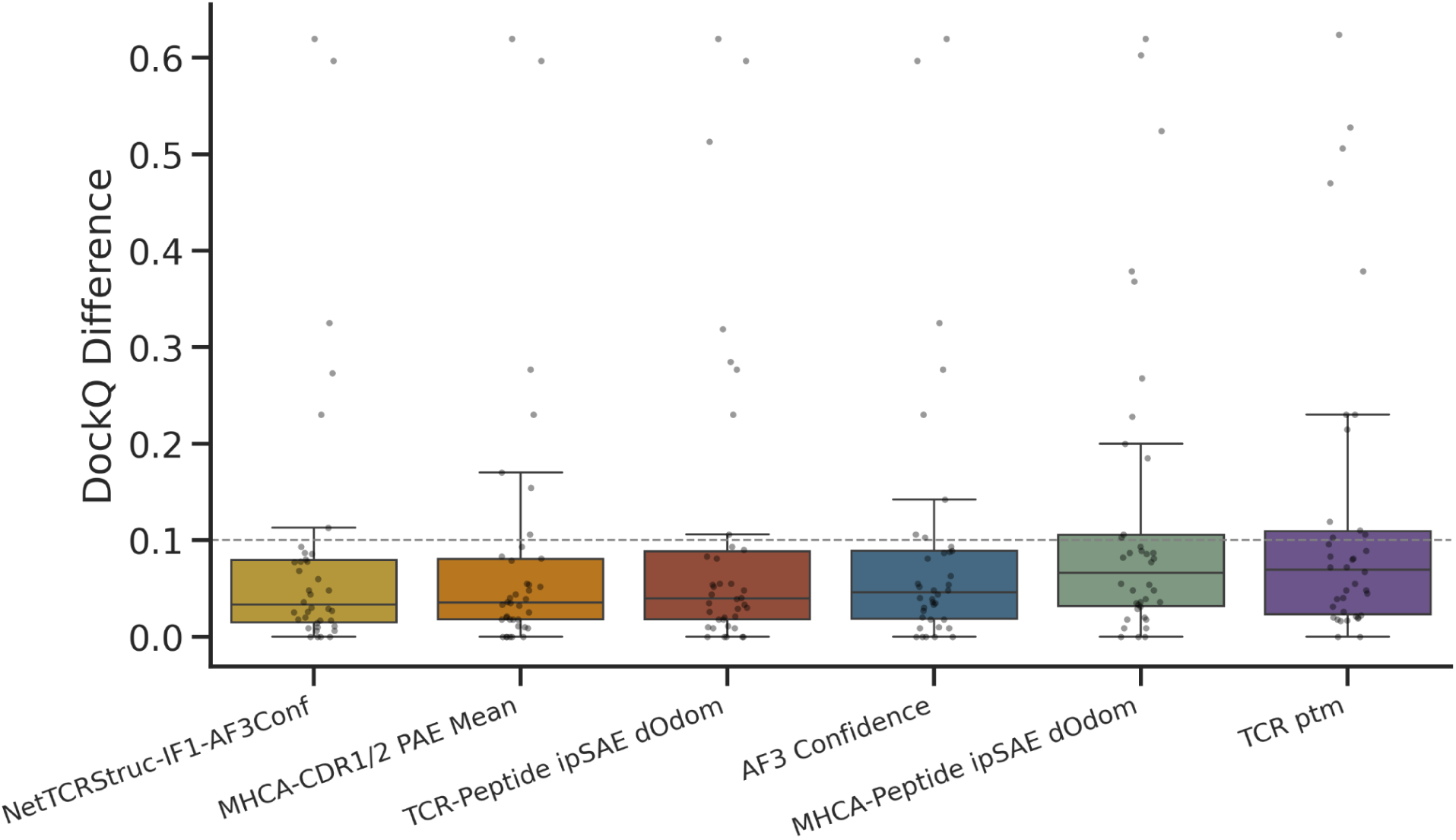
DockQ difference between Oracle Selection and AF3-based metric selection, per datapoint (n=38). Representative metrics from 10 best-performing (NetTCRStruct-IF1-AF3Conf, MHCA-CDR1/2 PAE Mean, TCR-Peptide ipSAE d0dom), medium-performing (AF3 Confidence) and worst-performing (MHCA-Peptide ipSAE d0dom, TCR pTM) groups are shown. Metrics are ordered by median DockQ difference, from lowest to highest.

**Supplementary Figure 5:**
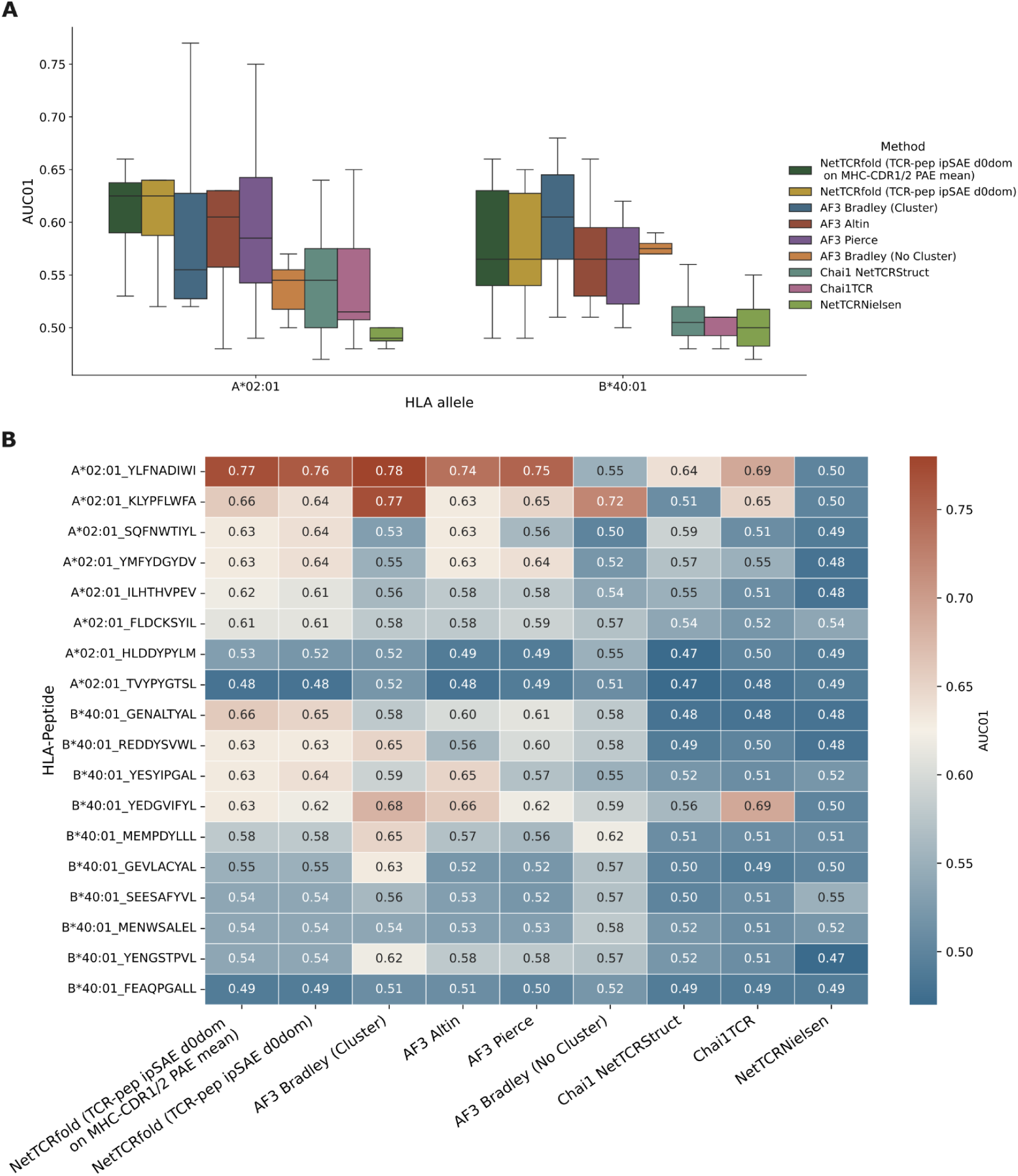
Per-allele and per-peptide breakdown of specificity prediction performance. A) AUC0.1 per HLA allele (HLA-A02:01, HLA-B40:01) for our pipeline and all evaluated Immrep2025 methods. B) AUC0.1 per individual peptide, across all 18 private peptides, for our pipeline and all evaluated Immrep2025 methods.

**Supplementary Figure 6:**
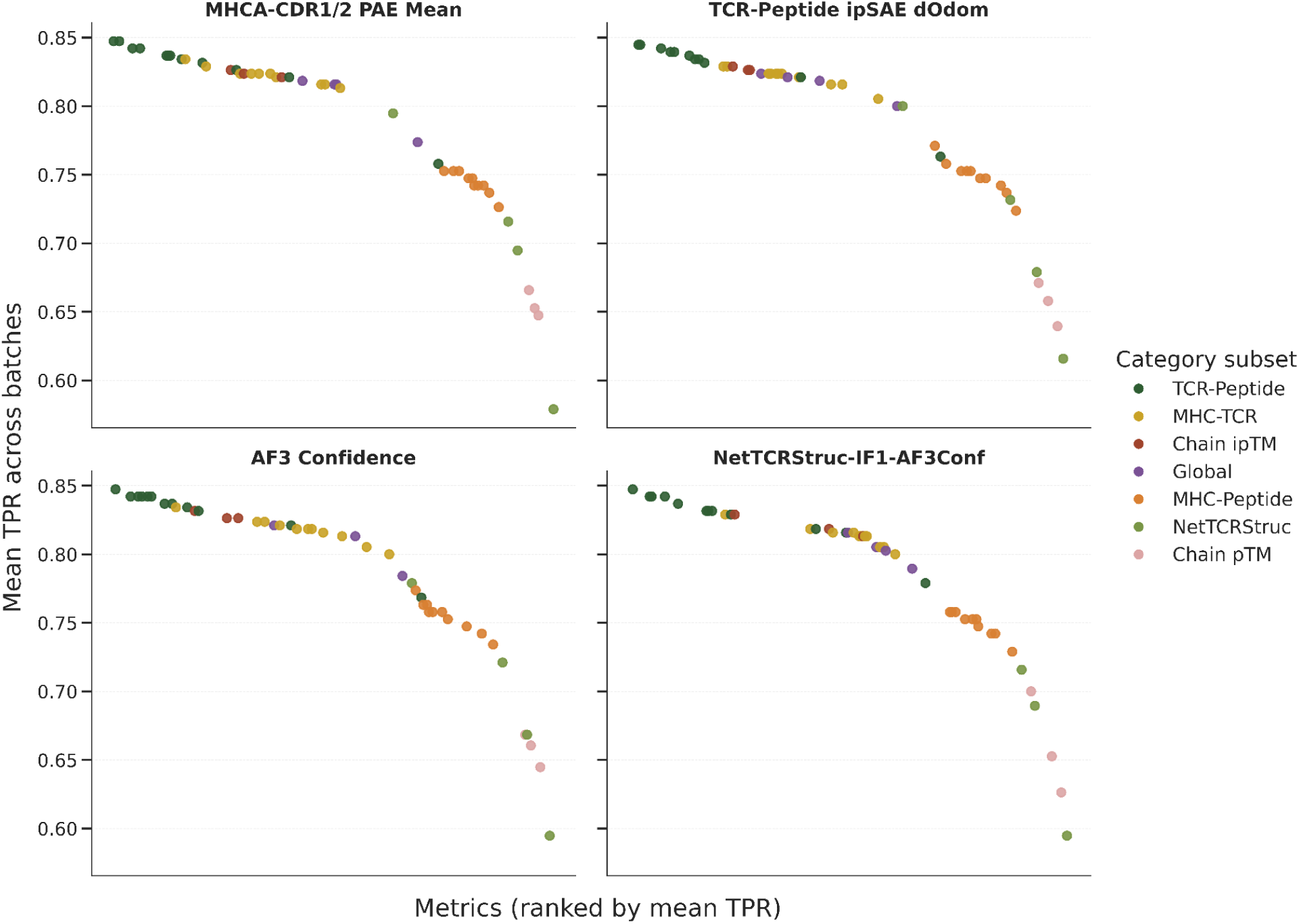
Performance of candidate metrics for target-selection with NetTCRstruc metrics included. For each model-selection metric (facets), performance is evaluated as the mean TPR across the 38 batches, using each candidate metric for target selection. Metrics are ranked by mean TPR. Each of the four facets shows the same set of target-selection metrics, with each point representing one metric, colored by its category as indicated in the legend. TCR-peptide, MHC-TCR and MHC-peptide categories group metrics by the interface they capture. Chain ipTM and chain pTM categories group single-chain metrics (TCR, peptide or MHC ipTM or pTM). Global metrics (AF confidence, ipTM and pTM) consider the complex as a whole. NetTCRStruc category contains the scoring metrics derived from NetTCRstruc.

**Supplementary Figure 7:**
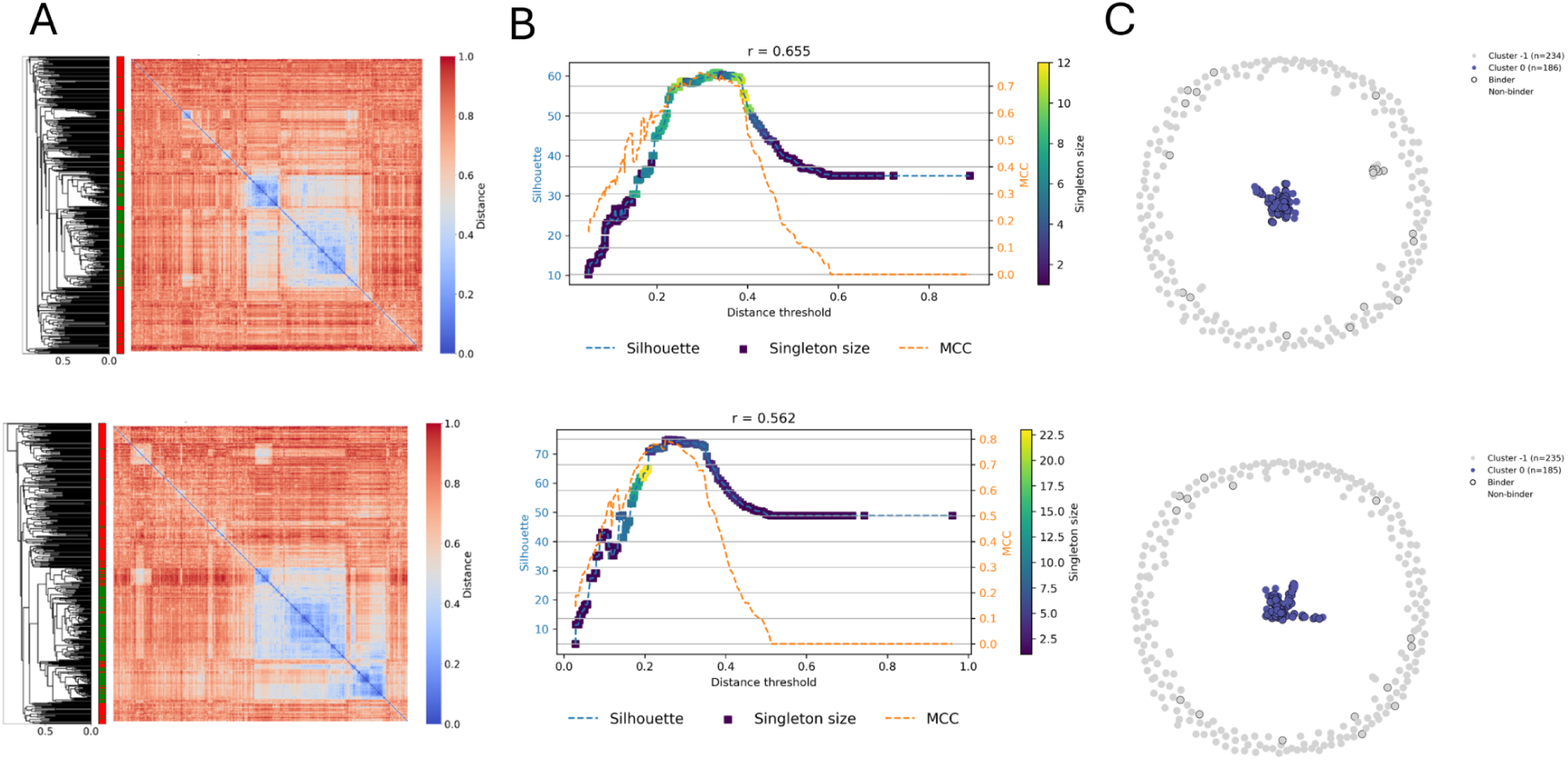
Overview of the sequence-based denoising pipeline, TCRdenoise, applied to the peptide-specific TCR repertoire for the YLQPRTFLL peptide from Messemaker et al. ^19^. The pipeline takes as input a set of TCR. A) In the first step, a matrix of pairwise TCR distances is calculated using a given similarity metric (here shown in the upper row for TCRbase ^39,40^ and in the lower row TCRdist3 ^41,42^). Smaller distances indicate higher sequence similarity. The side annotation indicates validated binders (green) and non-validated binders (red), and the dendrogram shows the hierarchical clustering used to order the heatmap. B) Next, the optimal cluster solution is identified using a refined Silhouette score. This refinement introduces a pseudo-cluster consisting of TCRs from healthy control negatives, singletons or clusters below a minimum cluster size threshold. This pseudo-cluster is included when calculating individual TCR Silhouette scores but excluded from the global Silhouette score, which is computed as the sum of the remaining TCR silhouette scores. To determine the optimal minimum cluster size threshold, a parameter search is performed over all possible cluster size thresholds, where clusters with sizes less than or equal to the threshold are assigned to the pseudo-cluster. The threshold yielding the highest global Silhouette score is selected. The Silhouette score curve is then coloured according to the selected minimum cluster size threshold. In cases where the TCRs contain a true label, the Matthews correlation coefficient (MCC) across the range of distance thresholds is also reported, and the Pearson correlation coefficient (PCC) between the Silhouette and MCC curves is shown above each plot. Here, showing a close to perfect co-location of the cluster solution with the max Silhouette score and the optimal MCC value C) Finally, the clustering solution at the distance threshold corresponding to the highest silhouette score is shown. Grey clusters represent the pseudo-cluster, whereas coloured clusters represent likely binders. In case of annotated data, TCRs outlined in black correspond to validated binders, while TCRs without an outline correspond to non-validated binders. Data and TCR annotations used here are from Messemaker et al. ^19^.

**Supplementary Figure 8:**
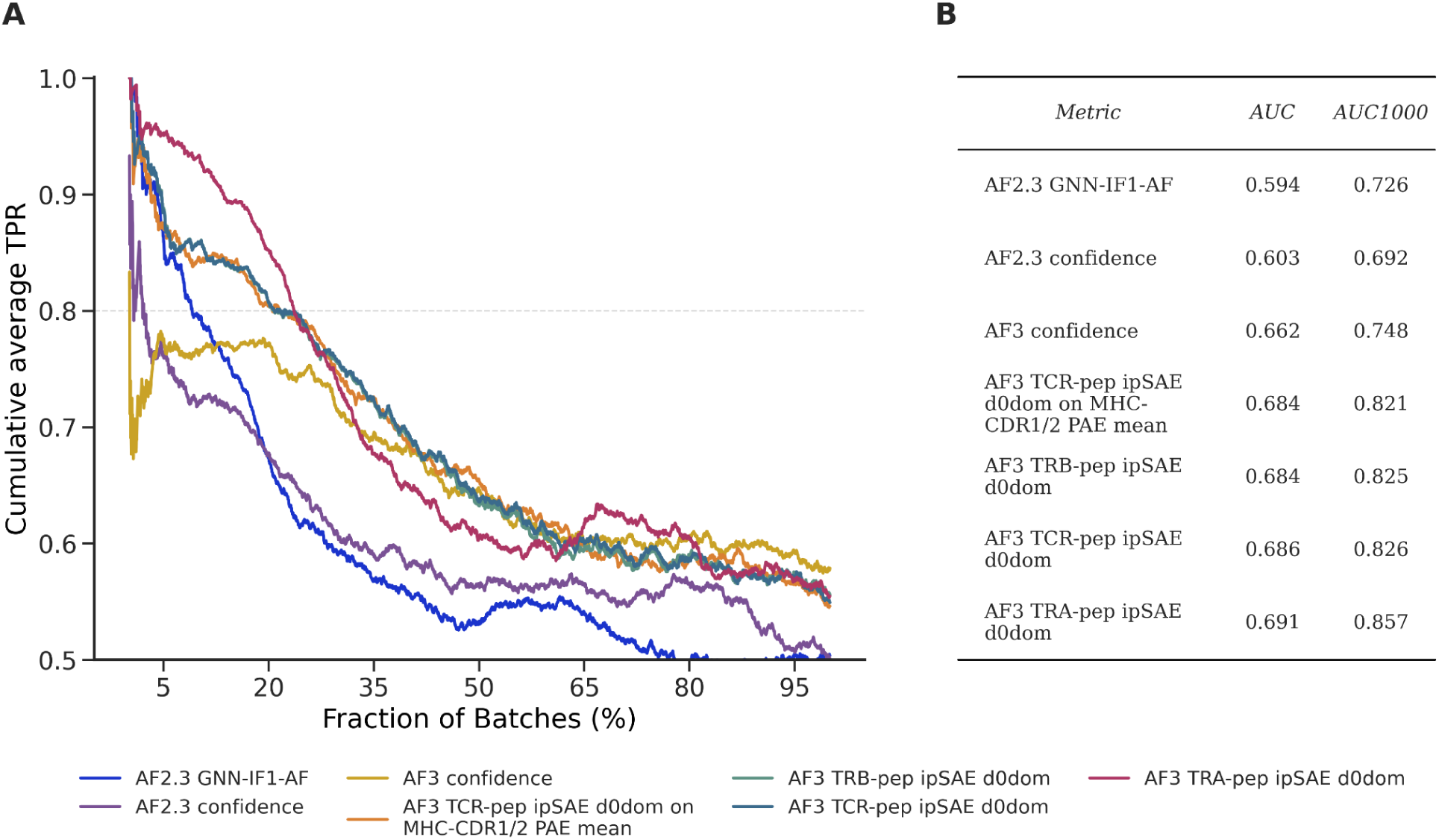
Specificity prediction performance across methods and metrics. A) Windowed cumulative average TPR across batches, sorted by maximal intra-batch score for each metric and displayed as a fraction of total batches, with a 500-batch rolling window. The legend indicates the metric used for target selection. If a different metric was used for model-selection, it is indicated by appending “on {metric to model selection}” to the metric name (see Methods for metric definitions). B) Table showing the AUC across all batches (AUC) and AUC restricted to the first 1,000 batches (AUC1000) for each metric in panel A.

